# Origin of Schooling and Collective Environmental Adaptation in Zebrafish

**DOI:** 10.64898/2026.07.01.732092

**Authors:** Mingyang Chen, Peishi Wang, Bo Li

**Author notes:** **Correspondence and requests for materials** should be addressed to B. L.

## Abstract

Zebrafish exhibit intricate schooling behaviors when swimming as a group. Such collective motion serves profound ecological functions and continuously inspires the design of highly coordinated artificial systems. Although the features and functions of schooling have been extensively studied, how this behavior originates over the course of individual development remains unknown, limiting a comprehensive understanding of its biological consequences. To address this gap, we developed a cross-scale, multi-modal experimental platform to capture zebrafish schooling and integrated AI-based algorithms to track fine-scale body posture and eye movements. We find that schooling emerges within a discrete developmental window, coinciding with coordinated changes in locomotor architecture and visual perceptual capacity. Specifically, structural remodeling of the caudal fin, enhanced muscle bundling, and an expanded visual perceptual range together provide the physical and sensory basis for the stabilization of polarized group movement. Network analyses under different representational frameworks reveal that the biological function of schooling is a collective group strategy for adapting to the external geometric environment. Our work provides a fundamental explanation of zebrafish schooling from a developmental perspective and elucidates a collective strategy that could inform the design of underwater robot arrays.

## Introduction

Collective behavior, coordinated local-interaction patterns [1], has independently evolved as an adaptive strategy that enhances fitness through energy savings, foraging, predation avoidance, and environmental responsiveness [2–10,12–16], with self-organizing principles underpinning ecological success and inspiring swarm intelligence and bio-inspired engineering [18–21]. Schooling, highly coordinated fish locomotion with tight cohesion and strong polarization [11,17], is among the most organized vertebrate collective behaviors, lowering predation via dilution and rapid information transfer [22–25], reducing energy costs, and serving as immediate adaptation and long-term strategy under natural selection [28]. Fish schooling research now integrates behavioral description with studies of dynamics, development, ecology, and genetics [29–33,34,35,36–44,58]. Schooling has been linked to local interaction rules and information flow [31–33,35], sensory/locomotor development [34,36], environmental adaptation [22–25,37–42], hydrodynamic interactions [26,27], and genetic architecture [43–44].

Despite these advances, current frameworks largely assume that individuals already possess mature and stable sensorimotor capacities, forming ordered structures through fixed local interaction rules. Consequently, existing theory primarily explains how schooling is maintained and what adaptive benefits it provides, but offers limited mechanistic insight into how highly polarized structures emerge during development. In particular, it remains unclear whether schooling arises from the gradual strengthening of local interaction rules or from stage-specific transitions triggered by functional maturation and reorganization of information networks. Most models abstract interactions into fixed rules based on spatial proximity or relative orientation and rarely incorporate realistic perceptual constraints, such as visibility limits, directional biases, or occlusion effects, into analyses of collective topology. A more fundamental question, therefore, remains: is the emergence of highly polarized schooling simply the outcome of progressively strengthened local interactions, or does it depend on a developmental reconfiguration of sensorimotor function and collective information networks?

To address this, we use zebrafish and a developmental, cross-scale framework to study schooling formation and adaptive function. We recorded locomotor stability and group polarization across developmental stages, testing schooling behaviors as a function of age. Combining sensory-behavioral assays with automated tracking, we build interaction network models that incorporate visibility and directional constraints, integrating perceptual limits into collective topology. By imposing environmental turning perturbations on mature groups, we examine dynamic network responses and adaptive reorganization. Our results show that schooling is not a gradual reinforcement of local rules but a stage-specific transition (**Fig. 1**) accompanied by sensorimotor maturation (**Fig. 2**) and vision establishment (**Fig. 3**). Mature networks confer adaptive flexibility in response to environmental cue (**Fig. 4**). Linking development characterization, multisensory remodeling assay, collective dynamics, and information network, this study elucidates the origin of fish schooling and its role in environmental adaptation.

**Figure 1.**
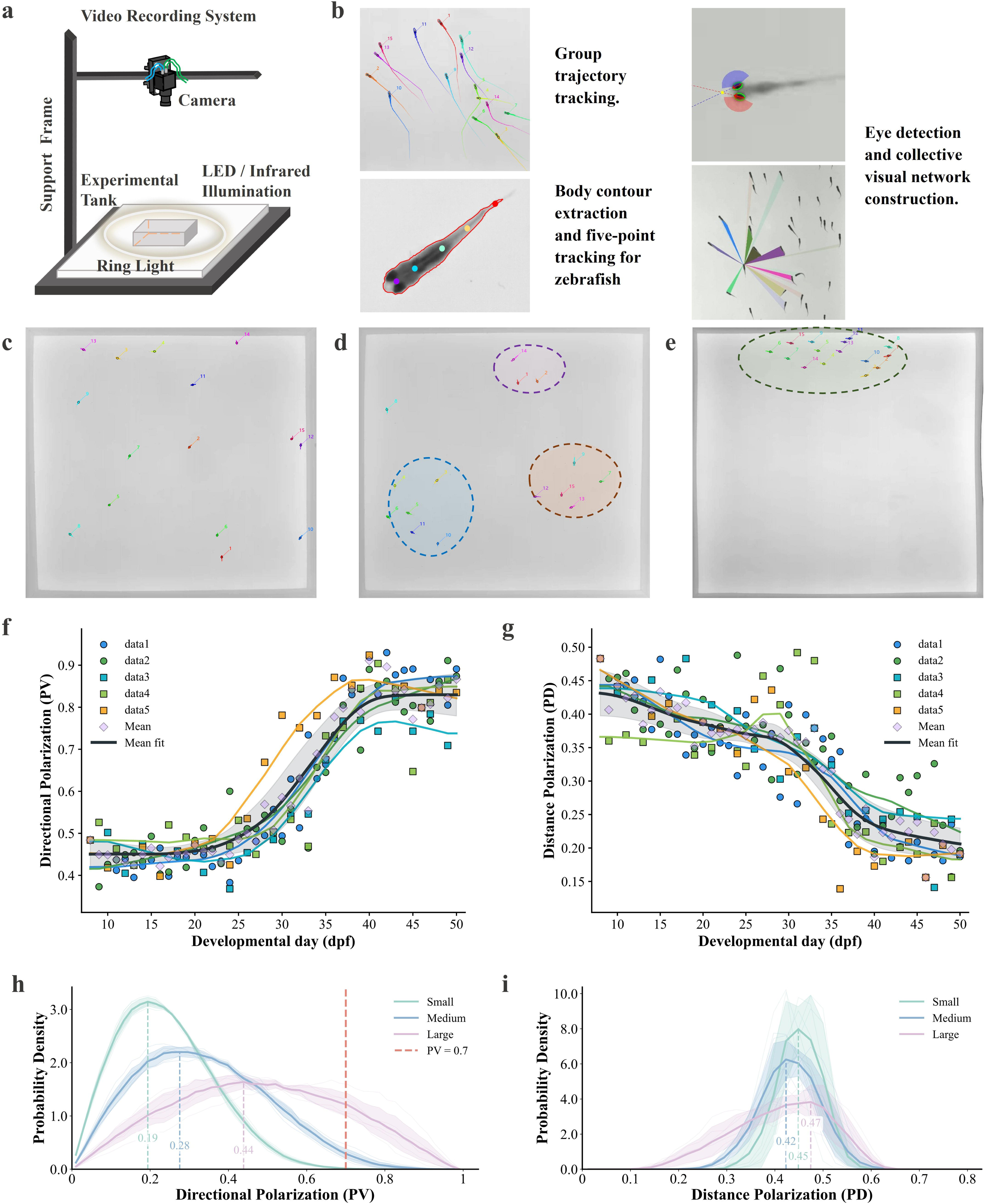
Experimental system for zebrafish collective behavior and its overall changes across developmental stages. **a,** Schematic of the video acquisition system for zebrafish collective behavior experiments. The experimental setup consists of an adjustable-height frame, a top-mounted industrial camera, a backlight source at the bottom of the tank, and a ring light surrounding the tank sides. The lighting system can switch between visible and infrared light [46,47,48] to accommodate illumination and behavioral recording requirements under different experimental conditions. **b,** Automated analysis pipeline for zebrafish collective behavior and construction of the visual network. The panel shows group trajectory tracking, where individual zebrafish are identified and trajectories are displayed in different colors with unique IDs. It also shows body contour extraction and five-point body midline labeling, with colored key points indicating body posture and a red outline representing the body contour. Eye detection is illustrated by the left and right eyes, head orientation, and visual field sectors, with yellow markers indicating the convergence points of visual directions. The panel further illustrates collective visual network construction based on visual distance and occlusion between individuals, where rays of different colors indicate other fish perceived within an individual’s visual field and their respective posture contours. **c-e,** Overall spatial distribution of zebrafish groups in the tank across developmental stages. (c) Zebrafish at about 10 days post-fertilization. Individuals are relatively dispersed in the tank with no obvious group structure. (d) Zebrafish at about 30 days post-fertilization. Several small groups begin to form, with representative groups highlighted by blue, purple, and orange circles. (e) Zebrafish at about 45 days post-fertilization. Individuals are highly aggregated, forming a stable group distribution with small inter-individual distances and overall aligned movement direction; the aggregated group is highlighted by a green circle. **f-g,** Quantitative characterization of zebrafish collective behavior during development. Long-term videos were recorded for zebrafish from 10 to 50 days post-fertilization, yielding five independent experimental datasets. The left panel (f) shows the change in group polarization (PV) across developmental stages; the right panel (g) shows the change in group positional dispersion (PD) across developmental stages. Colored dots represent raw data from different experiments, corresponding curves show fitted trends, the black curve represents the mean trend across all data, and dot shapes distinguish between two different tank conditions. **h-i,** Probability density distributions of zebrafish collective behavior over long timescales. Probability density distributions of PV (h) and PD (i) computed from long-duration recordings (about 1.5 h) of zebrafish groups at different developmental stages (Small, green; Medium, blue; Large, magenta). Light curves correspond to independent experimental replicates (n = 6), dark curves represent the averaged distributions, and shaded regions indicate variability across replicates.

**Figure 2.**
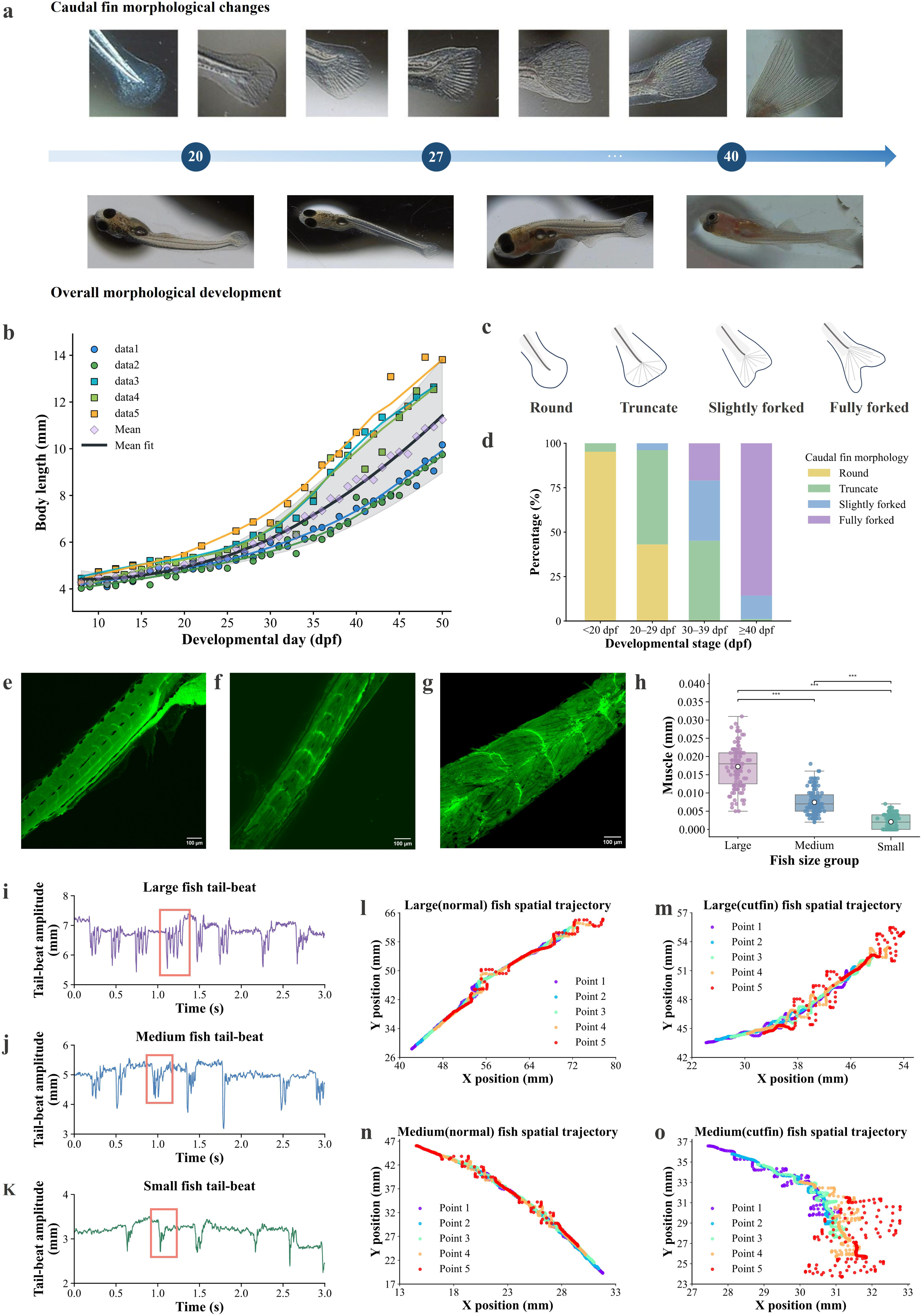
Developmental changes in zebrafish morphology and locomotor system. **a,** Schematic representation of key morphological features of zebrafish from 10 to 50 days post-fertilization (dpf). The timeline is indicated by arrows, with key developmental time points marked. Above the timeline, continuous changes in caudal fin morphology are shown; below, representative images of overall zebrafish morphology at corresponding developmental stages are displayed. **b,** Quantitative results of zebrafish body length across developmental stages. Measurements were obtained from five independent experimental datasets corresponding to the group behavior analysis in Fig. 1h and Fig. 1i. The x-axis represents age (days post-fertilization), and the y-axis represents body length (mm). Colored dots and shapes match those in Fig. 1h and i, indicating individual experiments; colored curves represent fitted trends for each dataset, and the black curve shows the mean trend across all data. **c,** Illustration of four key caudal fin morphologies: round, truncate, slightly forked, and fully forked. **d,** Distribution of caudal fin morphologies across developmental stages. The x-axis represents four developmental stages (<20 dpf, 20-29 dpf, 30-39 dpf, ≥40 dpf), different colors indicate the four fin types, and the y-axis shows the proportion of each morphology within the corresponding stage. **e-g,** Confocal imaging of muscle structure in zebrafish at different developmental stages. Representative images of small fish (e), medium fish (f), and large fish (g) are shown. All images were obtained from the transgenic line CZ26 at 10× magnification. **h,** Quantitative analysis of muscle bundle width across developmental stages (Samples lacking clearly identifiable muscle bundles were assigned a value of zero for quantitative analysis.). Each dot represents one measurement region sampled from multiple individuals (≥10 biologically independent fish per group). Statistical comparisons were performed using Welch’s t-test; asterisks denote significance levels. **i-k,** Illustration of tail-beat signals in zebrafish groups at different developmental stages, recorded using high-speed cameras. **l-o,** Comparison of tail-beat movements in individual zebrafish before and after caudal fin clipping. The left panels show trajectories of five key body points in large and medium fish before fin clipping; the right panels show corresponding body-point trajectories after fin clipping under the same conditions.

**Figure 3.**
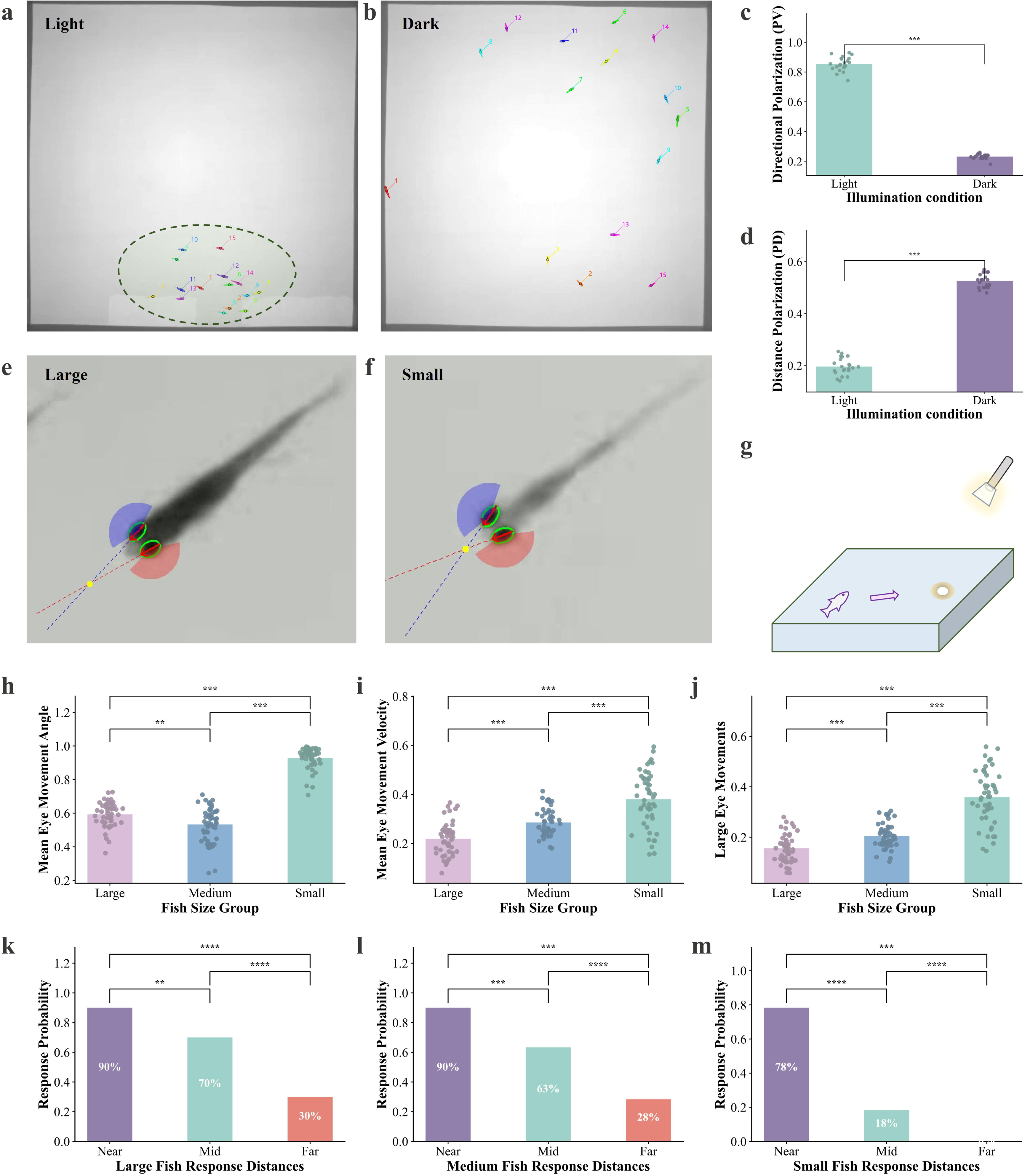
Group behavior and individual visual perception in zebrafish under different visual conditions. **a-b,** Representative spatial distributions of large-fish groups under visible light and darkness. (a) Group spatial distribution recorded under visible light. Different colors and numbers indicate individual fish, and the green circle highlights the overall group structure formed under visible light. (b) Group spatial distribution recorded under darkness using infrared bottom illumination [46,47,48]. Different colors and numbers indicate individual fish. **c-d,** Quantitative comparison of group behavior metrics under light and dark conditions. (c) Directional polarization (PV) of the group. (d) Distance polarization (PD) of the group. Each dot represents one independent experimental trial (one video recording). Statistical significance was assessed using Welch’s t-test; asterisks indicate significance levels. **e,** Schematic of eye-tracking and visual geometric model construction. Based on DeepLabCut [50] annotations of left and right eyes and head keypoints from Fig. 1d, ellipses were fitted to each eye to determine visual axis direction and major/minor axis parameters. The intersection of the extended visual axes determines binocular convergence (yellow marker). Lateral visual fields of about 150° were constructed for each eye based on visual axis orientation (left eye red, right eye blue). The panel shows results for large fish. **f,** Schematic of visual structure in small fish, showing closer binocular convergence. **g,** Schematic of the light-spot stimulus experiment. In a dim environment, brief light spots were presented at different distances to test behavioral responses of large, medium, and small fish. Response types included detectable behaviors (e.g., exploration or avoidance) and no obvious response **(Movie S5)**. **h-j,** Quantitative analysis of eye-tracking results comparing zebrafish at different developmental stages. **h,** Proportion of events with large eye-axis–body-midline angle (mean angle >7°). **i,** Proportion of large-amplitude eye movement angular velocity events (mean >200°/s). **j,** Proportion of events satisfying both criteria above. Each dot represents one fish (biological replicate). Comparisons across developmental stages were assessed using Welch’s t-test. **k-m,** Quantitative results of light-spot distance-response tests. Proportion of individuals showing detectable behavioral responses to light stimuli at close (0-5 cm), medium (5-10 cm), and far (>10 cm) distances across developmental stages. (k) Large fish; (l) Medium fish; (m) Small fish. Each data point represents the proportion of responding fish (n = 60 per stage). Statistical comparisons were performed using a two-proportion Z-test; asterisks indicate significance levels.

**Figure 4.**
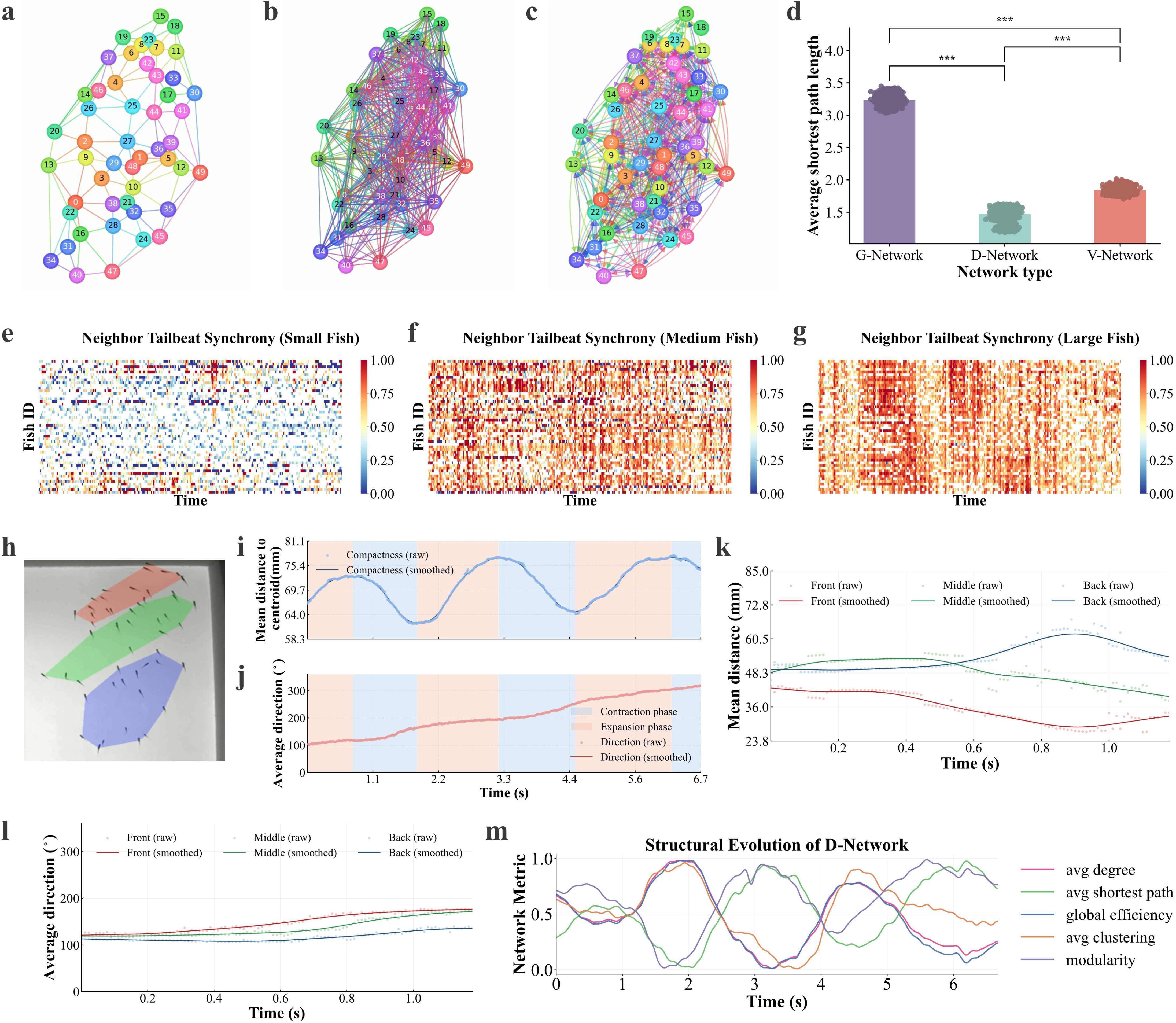
Group interaction networks under visual perception constraints and structural-environmental adaptation during group turning. **a-c,** Schematic of group interaction networks constructed from real experimental data (n = 50) under different interaction definitions. (a) Neighborhood network (G-Network) based on Delaunay triangulation. Undirected connections are established between any geometrically adjacent neighbors. (b) Distance-constrained network (D-Network) incorporating experimentally measured visual perceptual distance. Undirected connections are established between individuals within the effective perceptual range determined from the light-spot stimulus experiments in Fig. 3. (c) Visual network (V-Network) incorporating real visual perception and occlusion constraints. Using eye-tracking data to determine visual axis direction, experimentally measured visual distance, and individual body contours, ray-casting algorithms determine actual visible relationships. Connections are only established between mutually perceivable individuals. Due to directional visual perception, the network is directed, reflecting asymmetry in information perception. **d,** Quantitative comparison of three network structural metrics, showing mean shortest path length. Each dot represents one frame from the same video. Network metrics were computed for each frame using three different network construction methods. Pairwise comparisons of metric distributions across frames between methods were performed using Welch’s t-test. **e-g,** Tail-beat synchrony heatmaps under the V-Network. For each individual, the proportion of network neighbors tail-beat synchronized with it was calculated to quantify local interaction-level movement coordination. Color scale indicates synchrony intensity, with red for high synchrony and blue for low. (e) Small fish. (f) Medium fish. (g) Large fish. **h,** Schematic of spatial partitioning during group turning. According to each individual’s relative position along the group movement direction, the group is divided into front (red), middle (green), and rear (blue) regions, with shading indicating original trajectory data. **i-j,** Temporal changes in group structure and motion state. (i) Mean distance of individuals to the group centroid. (j) Mean overall movement direction of the group. Scatter points represent experimental data; curves represent fitted trends. Light blue and light orange shaded areas indicate contraction and expansion phases of the group envelope, respectively. **k-l,** Partitioned analysis corresponding to the first light-blue shaded interval in (i, j) during turning. Temporal changes in distance to centroid (k) and movement direction (l) are shown for front, middle, and rear individuals; curve colors correspond to partitions in (h). **m,** Dynamics of network metrics over time during interaction with the environment in a square tank, based on the D-Network. Displayed metrics include mean degree, mean shortest path length, global efficiency, mean clustering coefficient, and modularity.

## Results

### Establishment of Schooling Behavior with Age

Schooling is a highly coordinated form of collective motion in fish, characterized by spatial aggregation and directional polarization [11,17]. However, how this ordered behavior is progressively established during development remains poorly understood. During development, zebrafish undergo substantial changes that may underlie the emergence of schooling. To quantify this process, we performed long-term behavioral recordings of zebrafish groups across three developmental stages: small (<27 dpf), medium (27-39 dpf), and large (≥40 dpf), and, combined with automated trajectory tracking, analyzed group spatial organization and movement coordination **(Fig. 1a-b, Movie S1)**.

Groups at different developmental stages exhibited distinct spatial organization and movement patterns. In representative early-stage small fish (about 10 dpf), individuals were dispersed with large inter-individual distances and little directional consistency, resulting in disordered motion **(Fig. 1c)**. In medium fish (about 30 dpf), individuals began to form local clusters and exhibit emerging directional alignment, and the group displayed initial signs of local organization **(Fig. 1d)**. In large fish (about 45 dpf), individuals aggregated into a cohesive unit with reduced inter-individual distances and highly aligned movement directions, forming stable schooling **(Fig. 1e)**. These dynamics are also shown in **Movie S2**. Trajectory reconstruction further supported this transition: small fish exhibited disordered trajectories; medium fish showed emerging co-directional movement; and large fish displayed highly regular circular group trajectories, reflecting mature polarized collective motion **(Extended Data Fig. 1a-c)**.

To quantify the establishment of schooling, we calculated directional polarization (PV) and distance polarization (PD) [35,45]. PV remained low during early development, increased after about 27 dpf, and stabilized around 40 dpf **(Fig. 1f)**, while PD decreased progressively, with the most pronounced change between 27 and 40 dpf **(Fig. 1g)**. These trends were consistent across five independent experiments. Analysis of PV distributions revealed that highly polarized states (PV > 0.7) were nearly absent in small fish, appeared sporadically in medium fish, and became more frequent in large fish. Statistics over long-duration recordings (about 1.5 h per group) showed that the distributional patterns were consistent with the above results and varied with development **(Fig. 1h, i)**. Consistently, the fraction of time spent in highly coherent group states (PV > 0.7 and PD < 0.3) increased strongly in large fish but remained nearly absent in small fish **(Extended Data Fig. 1d)**. Mean swimming speed and acceleration calculated from representative short segments (20 s) increased systematically from small to large fish **(Extended Data Fig. 1e, f)**. In contrast, long-duration recordings (about 1.5 h) revealed probability density distributions of swimming speed and acceleration that shifted with development, with small fish exhibiting low, narrow, and sharp peaks, and medium and large fish showing broader and flatter distributions with higher values **(Extended Data Fig. 1g, h)**. In essence, zebrafish schooling is absent early and emerges within a defined window (around 27-39 dpf), after which stable collective behavior is established. This transition follows a clear developmental timescale and is therefore coupled to the maturation of certain organism body functions. In the next two sections, we uncovered the underlying mechanisms for these collective behavioral phenotypes. In the same developmental time window, the motor and visual systems undergo coordinated maturation, providing the physical basis for stable group polarization.

### Development Provides the Motion Ability for Schooling

**Figure 1** identifies a critical developmental window for schooling in zebrafish (around 27-39 dpf). Because stable polarized collective motion requires sustained and efficient propulsion, we asked whether locomotor maturation provides the physical basis for its emergence. We therefore systematically analyzed developmental changes in morphology and motor output in zebrafish from 10 to 50 dpf **(Fig. 2)**.

We first characterized overall body shape and caudal fin structure [36,51] across development using a standardized imaging system **(Extended Data Fig. 2a)** and summarized the evolution of key morphological features **(Fig. 2a)**. Caudal fin morphology exhibited a staged transition from rounded to truncate, then to slightly forked, and finally to fully forked forms. In parallel, body morphology changed progressively with development **(Fig. 2a)**. To quantify fin development, we classified fins into four categories (round, truncate, slightly forked, fully forked; **Fig. 2c**) and computed their proportions across stages **(Fig. 2d; Extended Data Fig. 2c)**. Round fins dominated before 20 dpf; truncate and slightly forked forms increased between 20-29 dpf; morphology diversified between 30-39 dpf with rising proportions of forked fins; after 40 dpf, fins were predominantly fully forked. These results indicate that caudal fin structure undergoes a discrete remodeling window rather than continuous change. In contrast, body length increased smoothly with development [51], consistently across five experiments **(Fig. 2b)**, indicating that somatic growth and maturation of the propulsive organ are temporally decoupled.

At the level of muscle structure, we further analyzed trunk muscle morphology across developmental stages. Confocal imaging at 10× revealed that in representative small fish, muscle tissue remained relatively fine-grained, with cross-sectional bundles often weak or difficult to resolve, whereas in large fish, structural units within muscle cross-sections became clearly discernible and exhibited stable bundled organization **(Fig. 2e-g)**. At higher resolution (40×), recognizable muscle bundling became more apparent with development and appeared further strengthened in large fish **(Extended Data Fig. 2d-f)**. Correspondingly, quantitative analysis demonstrated that the cross-sectional width of muscle bundles increased significantly across developmental stages **(Fig. 2h)**, supporting a progressive maturation of the propulsion-related muscular system from small to large fish.

At the level of motor output, we compared tail-beat patterns across developmental stages **(Fig. 2i-k)**. Pose reconstruction showed that small fish primarily exhibited isolated tail beats dominated by postural adjustment, medium fish displayed emerging consecutive tail beats, and large fish showed consecutive tail beating as the dominant locomotor pattern. Such consecutive beats provide sustained thrust, marking mature swimming capacity. To test the contribution of the caudal fin and musculature to propulsion, we performed caudal fin clipping **(Extended Data Fig. 2b)**. In large fish, tail-beat rhythms remained relatively stable after clipping, with preserved forward gliding, whereas in medium fish, clipping impaired propulsion, producing dispersed tail-beat trajectories and pronounced head oscillations **(Fig. 2l-o)**. Consistently, group-level analysis showed that in large fish, polarization (PV) and positional dispersion (PD) were unchanged after clipping **(Movie S3)**, whereas in medium fish, PV decreased and PD increased, indicating loss of group coordination **(Extended Data Fig. 2g, h)**.

In brief, the zebrafish motor system undergoes coordinated maturation around 27-39 dpf, with concurrent caudal fin remodeling, muscle bundling, and establishment of consecutive tail-beat patterns supporting stable, efficient propulsion. This window closely overlaps with schooling formation, indicating that locomotor maturation is a necessary prerequisite for polarized collective motion and provides a physical basis for subsequent integration of social information. The fact that schooling behavior does not completely disappear after fin clipping suggests, however, that locomotor maturation is not the sole reason for the establishment of schooling.

### Vision is Required for Schooling

The results above show that zebrafish establish a mature motor system supporting sustained propulsion around 27-39 dpf. However, propulsive maturation alone cannot account for highly polarized group coordination, which also requires continuous perception and integration of conspecific position and motion. As vision is a primary source of spatial information in many group-living animals [36,52], we next tested whether visual input contributes to the maintenance of group structure and evaluated its role in zebrafish schooling.

First, we performed visual deprivation experiments comparing large-fish groups under visible light and darkness **(Fig. 3)**. Under visible light, groups were highly aggregated with small inter-individual distances and strong directional alignment, forming stable collective structures **(Fig. 3a)**. In darkness (infrared illumination [46,47,48]), groups rapidly disintegrated: individuals dispersed, inter-individual distances increased, and movement became disordered, resembling early-stage small fish **(Fig. 3b, Movie S4)**. Quantitative analysis confirmed this transition, with darkness reducing PV and increasing PD **(Fig. 3c-d)**, indicating that visual input is essential for maintaining spatial cohesion and coordinated collective motion.

To characterize the geometric constraints of vision on group behavior, we constructed a zebrafish visual perception model based on eye-tracking data **(Fig. 3e, f; Extended Data Fig. 3a)**. Using left and right visual axes, we defined lateral visual sectors as the effective sampling space of individuals. Based on previous studies reporting visual fields of about 130°-165° [56–60], we adopted an intermediate value of about 150° as a representative assumption. The intersection of visual axes was used to estimate binocular convergence depth as a measure of spatial integration scale. In large fish, convergence points were located farther in front of the body, indicating integration over wider spatial scales, whereas in small fish, convergence was closer, suggesting near-field–restricted visual integration **(Fig. 3e, f; Extended Data Fig. 3b)**.

To characterize visual processing during development, we first analyzed high-speed eye movement recordings (1000 fps) of zebrafish **(Extended Data Fig. 3d-g)**. Eye movement amplitude was quantified by the angle *θ* between the eye axis and body midline **(Extended Data Fig. 3d)**, together with angular velocity, allowing identification of fixation and active scanning states **(Extended Data Fig. 3e-g)**. When *θ* exceeded about 7° or angular velocity exceeded about 200°/*s*, individuals typically entered an active scanning or near-focus state; these values were therefore used as empirical thresholds for large-amplitude eye movements. Quantitative analysis revealed developmental differences in eye movement behavior **(Fig. 3h-j)**: small fish showed a higher proportion of large-amplitude eye movements, with higher angular velocity and larger eye-axis angles than large fish, indicating more frequent visual scanning, whereas large fish relied more on stable wide-field monitoring. We next assessed the effective visual perception range using a light-spot stimulus **(Fig. 3g, Movie S5)**. Large and medium fish showed high response rates at near **(0-5 cm)** and intermediate **(5-10 cm)** distances, whereas small fish responded reliably only to near stimuli, with little or no response at longer distances **(Fig. 3k-m; Extended Data Fig. 3c)**, indicating a developmental expansion of visual perception range.

Put succinctly, zebrafish rely on visual input to maintain polarized group structure. During development, the visual system expands perceptual range and reshapes visual sampling strategies. Together with motor maturation **(Fig. 2)**, these results indicate that schooling depends on the joint development of propulsion and vision, providing a basis for social information integration and interaction network formation.

### Hybrid Network Explains the Collective Strategy in Environmental Adaptation

Results in previous sections indicate that developing zebrafish establish a vision-sensorimotor collective framework for stable coordination, implying schooling transcends individual sensing. We therefore asked how multi-dimensional information organizes into group-level interaction patterns for environmental responses. We constructed three interaction networks: geometric (G) from spatial proximity, distance-based (D) incorporating perceptual range, and visually constrained (V) including direction and occlusion (**Fig. 4a–c; Extended Data Fig. 4a–c, Movie S6**). The G-network (Delaunay triangulation) captured only local neighborhoods, sparse and homogeneous (**Fig. 4a**). Incorporating perceptual distance yielded a dense D-network with increased connectivity (**Fig. 4b**); adding visual direction and occlusion reduced V-network connections (due to mismatch/blockage), creating directionality and asymmetry (**Fig. 4c**). Both D- and V-networks showed higher mean degree and shorter path length than the G-network, indicating enhanced connectivity and information transfer under visual constraints (**Fig. 4d; Extended Data Fig. 4e**). Density was highest for the D-network and intermediate for the V-network (**Extended Data Fig. 4g**), suggesting visual constraints prune redundant links while preserving global integration.

With both vision and configuration considered, V-network fully captures instantaneous organization across development **(Extended Data Fig. 5a–f)**. Networks in small fish were loose and lacked stable clustering **(Extended Data Fig. 5d)**, whereas medium fish showed emerging local subgroups **(Extended Data Fig. 5e)**, and large fish formed highly integrated global networks. Accordingly, large fish exhibited the shortest path length, while small fish showed a lower degree and density, indicating a limited interaction range; medium fish were intermediate **(Extended Data Fig. 5j-l)**. A weighted visual network (W-network) incorporating distance decay and visual contour proportion further provided a continuous measure of interaction strength **(Extended Data Fig. 5g-i)**. Tail-beat events were extracted from midline curvature **(Extended Data Fig. 4d, f)**, and neighborhood synchrony was quantified as the fraction of synchronized neighbors, generating spatial maps **(Fig. 4e-g)**. Synchrony was low and fragmented in small fish, but increased in medium and large fish, forming coherent local patterns, matching the key developmental window for schooling (**Fig. 1f, g**), as well as the alteration of tail-beat modes (**Fig. 2i-k**). Together, these results indicate that visually constrained networks support stable motor coordination.

To elucidate how different networks reflect the group strategy during environmental adaptation, we considered coordinated turning as a representative behavioral paradigm because it requires rapid group reconfiguration, providing an ideal context to evaluate the functional relevance of interaction-based representations. We divided individuals into front, middle, and rear positions according to their relative location along the group movement direction (**Fig. 4h**). During turning in a square arena, the group envelope underwent periodic contraction and expansion accompanied by changes in movement direction (**Fig. 4i, j**). Front individuals moved toward the group centroid first, followed sequentially by middle and rear individuals (**Fig. 4k**). Heading changes followed the same front-to-rear sequence (**Fig. 4l**), indicating that turning arises from orderly spatiotemporal propagation of positional and directional adjustments across the group. Similar envelope contraction preceding reorientation was observed in a rectangular arena (**Extended Data Fig. 6c-g**) and during group merging (**Extended Data Fig. 6h-j**), suggesting a general collective strategy for environmental adaptation.

The D-network best captures the group strategy during turning. In a square arena, group turning involved contraction and expansion of the group envelope – the oscillation of mean distance in **Fig. 4i**. Network metrics were strongly coupled to this modulation: during contraction, mean shortest path length and modularity decreased, whereas mean degree, clustering coefficient, and global efficiency increased; opposite trends occurred during expansion **(Fig. 4m)**. These results indicate that turning involves not only geometric compression but also functional network reconfiguration into a highly connected state. The V-network showed higher-frequency fluctuations **(Extended Data Fig. 6a)**, implying rapid perceptual updates at the individual level and increased sensitivity to fine-scale motion variability. In contrast, the D-network provides a more coarse-grained representation of interaction structure by filtering out short-timescale fluctuations while preserving distance-constrained coordination patterns, thereby capturing the periodic reorganization associated with turning dynamics. The G-network failed to capture this periodic reorganization **(Extended Data Fig. 6b)**.

Taken together, zebrafish form a hybrid interaction network under visual and spatial constraints that integrates local perception at the group scale. This network is reconfigured prior to environmental cues, optimizing connectivity and information transfer efficiency to support rapid collective adjustment. Different representations (G-network, D-network, V-network) of this multidimensional networks captures different aspects of the schooling behaviors (tail beat synchrony, group turning). Using this network-dynamical framework, we show that schooling arises not only from local rules but from vision-guided network organization enabling predictive adaptation..

## Discussion

Collective behaviour is often interpreted as a self-organized process arising from simple local interaction rules, in which alignment and aggregation reflect instantaneous neighbour responses. Here, adopting a developmental perspective, we dissect the ontogeny of zebrafish schooling and propose an integrative framework linking sensorimotor maturation to the emergence of collective network structure. Highly polarized schooling does not increase gradually with growth but instead emerges abruptly within a defined developmental window (**Fig. 1**). Locomotor maturation supplies the biomechanical substrate for stable propulsion (**Fig. 2**); visual integration expands and restructures the scale of social interaction (**Fig. 3**); and perceptually constrained networks embed individual capacities into collective information architecture, allowing schooling to emerge and reorganize efficiently under environmental challenge (**Fig. 4**). Together, these findings establish a mechanistic linking between individual developments to collective adaptation. Collective intelligence thus emerges not as an abstract property detached from ontogeny, but as the coordinated integration of sensorimotor capacity and information organization across development.

Our findings emphasizes the complexity of collective behaviors. With the course of development, the biological functions of living organisms systematically mature -for example, sensorimotor and perceptual capacities in this particular work – leading to information coupling at multiple representations. Importantly, this prerequisite-based organization has implications for collective artificial systems. Rather than relying solely on fixed local interaction rules, swarm intelligence can be improved by explicitly incorporating sensing, interaction constraints, and even brain activities into system design, so that coordination emerges through distributed sensing, adaptive reweighting of interactions, and structured integration of spatial information under changing environmental conditions [63,64]. From an ecological and evolutionary perspective, our findings suggest that the timing of sensorimotor maturation can constrain the onset and robustness of collective behavior, thereby shaping the emergence and stability of coordinated group function under environmental variability. This links developmental processes to collective performance in natural systems and highlights how developmental constraints influence adaptive flexibility in spatially structured environments [65,66].

## Online Methods

### Fish breeding

Zebrafish used in this study included wild-type (WT) and muscle transgenic (CZ26) strains. Adult fish were maintained in culture tanks equipped with recirculating water, oxygen supply, and temperature control; water temperature was maintained at about 26°C, the photoperiod was 14 h light / 10 h dark per day, and fish were fed fresh brine shrimp twice daily. All experimental larvae were bred and reared in the laboratory. Adults designated for breeding were separated by sex for at least two weeks prior to mating. On the night before spawning, zebrafish were transferred to spawning tanks for acclimation; the following morning, partitions were removed to allow natural mating, and fertilized eggs were collected. The day of fertilization was designated as 1 dpf (days post-fertilization). Embryos were incubated in still water at an appropriate temperature, with daily water changes. Larvae began feeding at about 6 dpf with starter feed, which was gradually adjusted according to developmental stage. At 16 dpf, larvae were transferred to the recirculating system and maintained under low-rate drip flow.

### Experimental system and video taking

The video imaging system **(Fig. 1a)** and the separate still-image acquisition setup **(Extended Data Fig. 2a)** were custom-built. The video arena was handmade from white polypropylene (PP) sheets and PP adhesive, with a bottom backlight and a surrounding ring of side lights to provide even and stable illumination. Arena dimensions were adjusted according to zebrafish body length at different developmental stages to ensure sufficient space for free swimming. Square arenas ranged from 10 × 10 cm to 50 × 50 cm, while the rectangular arena measured 119 × 60 cm. All experiments were conducted in a quiet environment, allowing fish adequate acclimation prior to filming, and ambient temperature was kept constant and suitable during recording. The imaging setup included an adjustable-height mount, industrial cameras (MindVision, MV-XGC1207GC/GM, up to 4090 × 3072 resolution, 30-90 fps), and a high-speed camera (SSZN, SH6-504-M-80, 250-1000 fps), combined with bottom backlighting and lateral ring lighting for multi-scale behavioral recordings. For visible-light experiments, soft bottom backlight with lateral ring fill was used; for dark-field behavior, only the infrared bottom light (Lighting & Optics Tech Specialist, LTS-3FT700700,850nm) [46,47,48] was turned on.

For developmental group behavior experiments, each group consisted of 15 zebrafish. All individuals were derived from the same batch of fertilized eggs, but different fish were used on each day to avoid repeated sampling. Arena size was adjusted according to body length at each developmental stage to ensure sufficient space for free swimming. For data1 and data2 in **(Fig. 1h-i)**, square arenas scaled by age from 10 to 40 cm were used, whereas data3-data5 employed a combination of wider 20 cm and 40 cm square arenas to minimize potential effects of spatial constraints on group structure. All behavioral experiments were conducted under visible light and recorded continuously at 30 fps with an industrial camera, with each group filmed for at least 1.5h without intervention. Data for group polarization (PV) and spatial dispersion (PD) were extracted from these long-duration recordings. For each video, the continuous 20s segment with the highest PV was automatically identified, and PD, as well as mean individual speed and acceleration, were computed for that segment. Additionally, over the full 1.5h time series, the proportion of frames satisfying both PV > 0.7 and PD < 0.3 was calculated to quantify the frequency of high-consistency group states across developmental stages. After behavioral recording, individual body morphology and caudal fin shape were documented. Morphological imaging **(Fig. 2a-d; Extended Data Fig. 2c)** was performed in the separate still-image setup, with a scale placed in the dish for body length calibration; body length measurements were conducted using ImageJ **(Fig. 2b)**.

Group interaction network construction (with arena size adjusted according to fish body size), partial eye movement statistics, and environment-response data **(Fig. 3e-f**, **Fig. 3h-j**, **Fig. 4; Extended Data Fig. 4, Extended Data Fig. 5, Extended Data Fig. 6a-b)** were based on data acquired by the industrial camera at 90 fps. Each experiment used 50 zebrafish per group under a unified imaging protocol to ensure comparability of network analyses across developmental stages. Environment-response experiments shown in **Fig. 4h-m** and **Extended Data Fig. 6a-b** were primarily conducted in the 50*cm* × 50*cm* square arena; validation experiments at larger spatial scale **(Extended Data Fig. 6c-j)** used the 119*cm* × 60*cm* arena and were recorded at 30 fps. The high-speed camera was employed to resolve fine kinematic features and eye movements: group-level individual pose dynamics **(Fig. 2i-k)** were recorded at 500 fps, caudal fin clipping experiments **(Fig. 2l-o)** at 250 fps, and a representative single-fish eye movement sequence was acquired at 1000 fps to characterize rapid eye movements and define empirical thresholds for large-amplitude eye movement events **(Extended Data Fig. 3d-g)**. Visual deprivation experiments **(Fig. 3a-b)** were implemented by switching between visible light and infrared bottom illumination [46,47,48], with all other environmental parameters kept constant.

### AI based poise and eye movement tracking

Individual identity, position tracking, contour fitting, and midline extraction in zebrafish groups were done with idtracker.ai [35,45,49] **(Fig. 1b**, **Fig. 1e-i, Figure S1, etc.)** and served as the basis for subsequent group interaction network construction **(Fig. 4a-g, Extended Data Fig. 4, Extended Data Fig. 5)**, environmental adaptation analysis **(Fig. 4h-m, Extended Data Fig. 6)**, and behavioral dynamics quantification **(Extended Data Fig. 1)**. The algorithm allows long-term stable tracking of multiple individuals without physical tags and outputs the spatial position and body contour of each fish in every frame.

Individual fine pose keypoint tracking (five body key points plus eye and head key points; **Fig. 1c**, **Fig. 2l-o**, etc.) was done with DeepLabCut [50] **(Figure S2)**. For fine body pose in the group **(Fig. 2i-k)** and eye movement tracking **(Fig. 3e-f**, **Fig. 3h-j, Extended Data Fig. 5b-i)**, a hierarchical pipeline was used to combine group-scale identity consistency with single-fish high-resolution pose. Specifically, idtracker.ai [35,45,49] was first used to extract trajectories of each fish in the group and crop local video clips at about 2× body length around the center position; DeepLabCut [50] was then run on each single-fish video to obtain stable keypoint coordinates.

The eye movement tracking pipeline in the group is shown in **Extended Data Fig. 3a** and **Figure S3**. In brief, idtracker.ai [35,45,49] was used to extract individual trajectories and generate corresponding single-fish video clips; DeepLabCut [50] was then used to automatically label left/right eye and head keypoints. On this basis, Python with OpenCV was used to automatically extract eye shape and orientation. For each frame, a local region centered on the eye key point was cropped and converted to grayscale; adaptive (or fixed) thresholding was used to obtain the eye contour, and under an area constraint the largest connected component was selected and fit with an ellipse using fitEllipse. The ellipse major axis was taken as the eye’s main orientation axis. To unify orientation, the eye-facing side was determined by comparing the distance from the two endpoints of the major axis to the head position, yielding per-frame eye center coordinates, major axis length, and orientation angle.

Individual binocular visual field sectors were built from left and right eye axis information, and their convergence region was computed to estimate instantaneous visual attention direction. For group-level network analysis, this visual geometry was mapped back to the original group coordinate system and, combined with individual pose (contour fitting from idtracker.ai [35,45,49]) and visual distance constraints **(Fig. 3k-m)**, a ray-casting algorithm was used to determine which neighbor individuals were actually within the visible range at each time, thereby excluding occluded targets and constructing the interaction network under real visual perception constraints **(Fig. 4c, Extended Data Fig. 5d-i)**.

### Slicing and microscopy

CZ26 transgenic zebrafish were collected in batches according to developmental stage. Samples were fixed in 4% paraformaldehyde (PFA) at room temperature for 6-8 h, then washed three times in PBS. Fixed tissue was transferred to 20% sucrose and kept at 4°C overnight until fully sunk for dehydration and cryoprotection. Before embedding, each fish was placed in a small mold containing embedding medium and pre-equilibrated at 4°C for about1 h; another mold was pre-filled with embedding medium at the bottom and frozen at −20°C until solid. The sample was then transferred into the frozen mold for embedding and fully frozen at −20°C.

Serial sectioning was done on a cryostat (epredia, HM525 NX); section thickness was 20 – 40 *μm* and sections were mounted on slides. After sectioning, slides were left at room temperature for at least 30 min to improve tissue adhesion. For long-term storage, sections could be kept at −20°C. Before imaging, slides were washed three times in PBS, mounting medium was applied, and coverslips were placed. Confocal imaging was done with 10× or 40× objectives. For early developmental stages with high overall transparency **(Fig. 2e-f)**, whole-mount confocal imaging (10×) was performed directly after PFA fixation. For larger, opaque individuals **(Fig. 2g)**, cryosectioning was followed by confocal imaging at 10×. For finer muscle structure, all samples in **Extended Data Fig. 2d-f**) were cryosectioned and imaged at 40× on the confocal microscope.

For muscle bundle width quantification **(Fig. 2h)**, measurements were performed on both CZ26 transgenic fish and immunofluorescence-stained transverse sections from wild-type zebrafish. Wild-type sections were permeabilized, blocked with 3% BSA, and stained using recombinant Anti-non-muscle Myosin IIB/MYH10 antibody (EPR22564-23), followed by goat anti-rabbit IgG H&L (Alexa Fluor® 594) secondary antibody and Alexa Fluor™ 488 Phalloidin. The two approaches yielded similar measurements, and data from both methods were included in the final analysis.

### Dynamical analysis

Group directional polarization (PV) and distance polarization (PD). To quantitatively characterize collective coordination during zebrafish group locomotion, we introduced two complementary metrics: directional polarization (PV) **(Fig. 1h**, **Fig. 3c, Extended Data Fig. 1d-g, Extended Data Fig. 2g-h)** and distance polarization (PD) **(Fig. 1i**, **Fig. 3d, Extended Data Fig. 2h).** These metrics describe, respectively, the coherence of movement direction at the group level and the spatial compactness among individuals, thereby capturing the global organizational state of the school during dynamic swimming. PV and PD were computed frame by frame and averaged over selected time windows to obtain group-level metrics for each interval. More details are offered in the Supplementary Information.

### Light stimulation

Light-stimulation experiments were conducted under dim ambient illumination. A uniform bottom backlight was provided beneath the arena, which was additionally covered with slightly light-transmitting black cloth and diffuser paper to minimize environmental interference. Zebrafish were placed in the arena at least 5 min prior to stimulation for acclimation. A brief white light spot was then presented above the water surface as the stimulus to evoke behavioral responses. Following stimulation, fish typically exhibited escape, exploratory, or no-response behaviors. After each trial, the distance between individual fish and the light spot was measured using ImageJ and used for subsequent statistical analysis **(Fig. 3g**, **Fig. 3k-m)**.

## Network construction and quantification

### G-Network

To characterize the instantaneous spatial organization among individuals in the group, an interaction network based on a geometric interaction network (G-network) was constructed. In this network, each fish was represented as a node, and edges were defined according to geometric neighborhood relationships. At each time frame, Delaunay triangulation [61] was applied to the two-dimensional positions of all individuals. Pairs of fish sharing a triangle edge were considered direct spatial neighbors and were connected by undirected links. Because the Delaunay network adaptively captures local nearest-neighbor structure without requiring an explicit distance threshold, it is widely used to describe spatial interaction topology in collective systems. The resulting G-network represents the instantaneous spatial connectivity of the group. By constructing the network independently for each frame, a time-resolved network sequence was obtained, enabling further analysis of the dynamic evolution of group structure.

### D-Network

To incorporate visual perceptual scale into group interaction structure, a distance-threshold–based geometric network (D-network) was constructed. In this network, each fish was represented as a node, and edges were established according to instantaneous spatial distance. Specifically, at each time frame, pairwise two-dimensional Euclidean distances were computed for all individuals, and an undirected edge was added between two nodes when their distance was smaller than a predefined threshold *d*. The distance threshold *d* was determined from the light-spot stimulation experiments **(Fig. 3k-m)**, which were used to estimate the maximal effective visual sensing range of zebrafish under the experimental conditions. Based on the distance-dependent behavioral responses to transient light stimuli at different developmental stages, the largest distance that still elicited a clear response was selected as the perceptual scale and converted into the trajectory coordinate system. In this study, the intermediate-distance threshold was used for medium and large fish, whereas the near-distance threshold was used for small fish. The D-network therefore represents connections between individuals that fall within mutual visual sensing range, reflecting potential interaction structure under explicit perceptual distance constraints. By introducing a physical length scale, this network becomes sensitive to changes in group density and spatial distribution, enabling characterization of group compactness, local aggregation, and overall spatial organization. Compared with the G-network, the D-network emphasizes direct spatial interactions constrained by visual accessibility, providing a complementary perspective for understanding how perceptual scale shapes collective structure.

### V-Network

On the basis of the D-network, a vision network [33,62] (V-network) was further constructed by introducing realistic visual constraints. This network jointly considers inter-individual distance, visual field range, and occlusion, and is used to characterize the directed perceptual structure formed under actual visual conditions. The V-network is a directed network, in which nodes represent individuals and edge direction indicates visual perception, i.e., from the observing fish to the target fish. Specifically, a directed connection was established between two individuals only when their instantaneous distance was smaller than the visual threshold *d*, the target individual lay within the effective visual field of the observer, and the target was not occluded by other individuals. Individual visual orientation was obtained from eye-movement tracking **(Extended Data Fig. 3a)** and combined with reconstructed spatial positions. Visibility relationships were then determined using a geometric ray-casting approach. Therefore, the V-network describes a directed visibility network under realistic visual constraints, reflecting potential information acquisition pathways among individuals in the group. Compared with the G-network and the D-network, the V-network further incorporates visual directionality and occlusion mechanisms, providing a closer approximation to actual sensory perception and offering a critical framework for investigating visually driven collective coordination behavior. More details are offered in the Supplementary Information.

## Acknowledgments

We thank Deepseek v2.2.1 for text polishing. This study was supported by taxpayers of China through National Natural Science Foundation of China (Distinguished Young Scholars Funding, Overseas; Young Scientists Fund - Category C) and Wuhan University (Talents Startup Funding).

## Author contributions

B.L. conceptualized the project. B.L. and M.Y.C. designed the research. M.Y.C. performed the experiment and analyzed the data with the help of P.S.W.. All authors discussed the results and wrote the paper.

## Competing interests

The authors declare no competing interests.

## Additional information

## Extended data

Is available for this paper after the main text.

## Supplementary information

Is available for this paper at a separate file.

## Reprints and permissions information

Are available at

## Data availability

The data and analysis code that support the findings of this study are submitted with the paper by separate files.

---

**Extended Data Figure 1.**
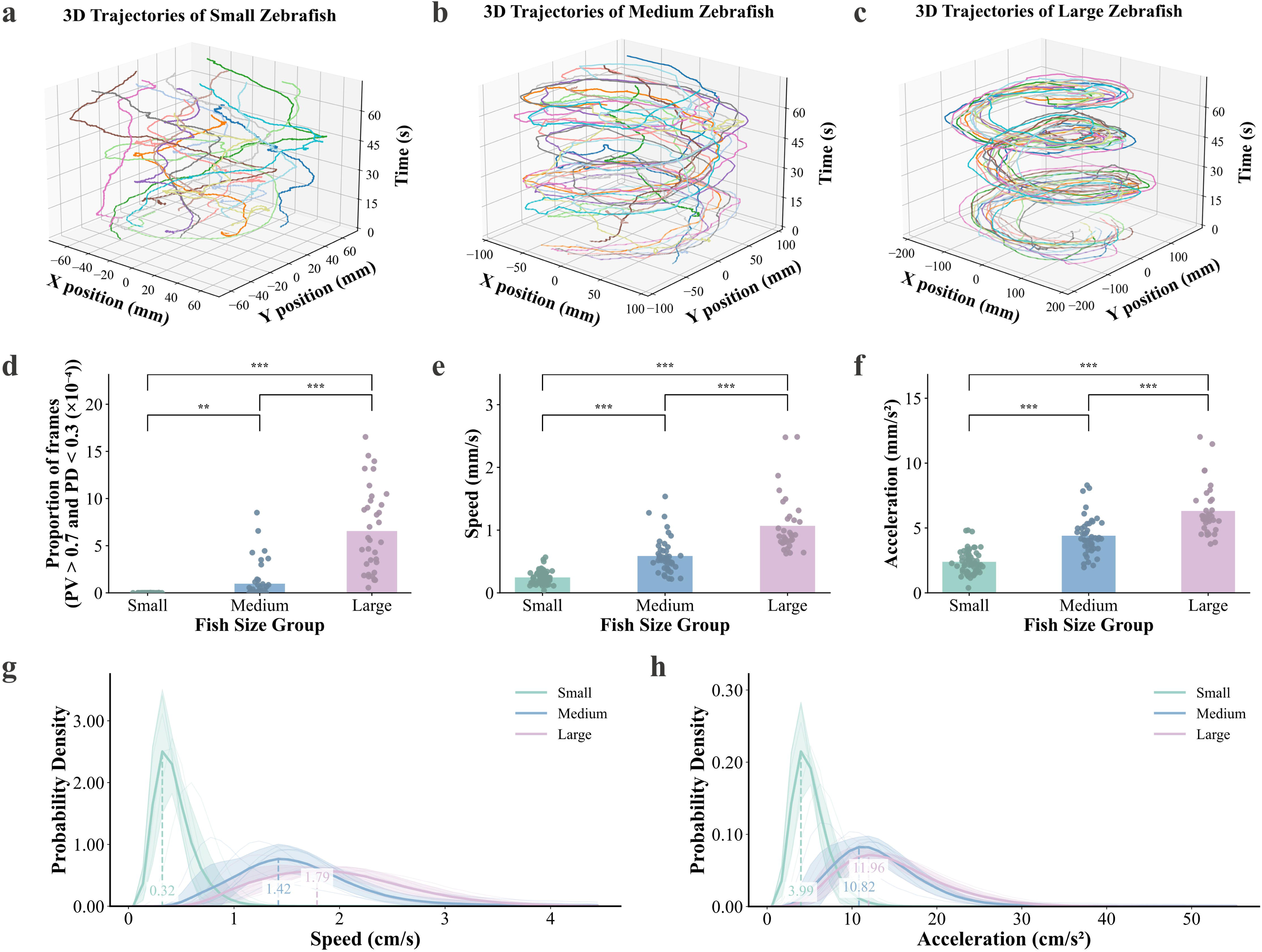
Collective locomotor structure and dynamics of zebrafish at different developmental stages. **a-c,** Schematic of 3D trajectories of 15 zebrafish in a single segment of group swimming video (X-Y spatial positions over time). Different colors indicate individual fish. (a) Small fish. (b) Medium fish. (c) Large fish. **d,** Proportion of high-consensus group states based on complete long-term recordings (about 1.5 h per group). Frames satisfying PV > 0.7 and positional dispersion index PD < 0.3 were counted and expressed as a fraction of total frames, grouped by developmental stage (small, medium, and large fish). Scatter points represent independent experimental samples. Comparisons between developmental stages were performed using Welch’s t-test. **e-f,** Average swimming speed (e) and average acceleration (f) calculated from 20 s representative video segments used for developmental analysis in Fig. 1. Scatter points represent independent experimental samples. Comparisons between developmental stages were performed using Welch’s t-test. **g-h,** Probability density distributions of locomotor dynamics computed from long-duration recordings (about 1.5 h) across developmental stages. (g) Swimming speed distributions. (h) Acceleration distributions. Colored curves represent different developmental stages (Small, Medium, Large; consistent color coding as in Fig. 1), with light curves corresponding to individual experimental replicates (n = 6) and dark curves representing averaged distributions. Shaded regions indicate variability across replicates.

**Extended Data Figure 2.**
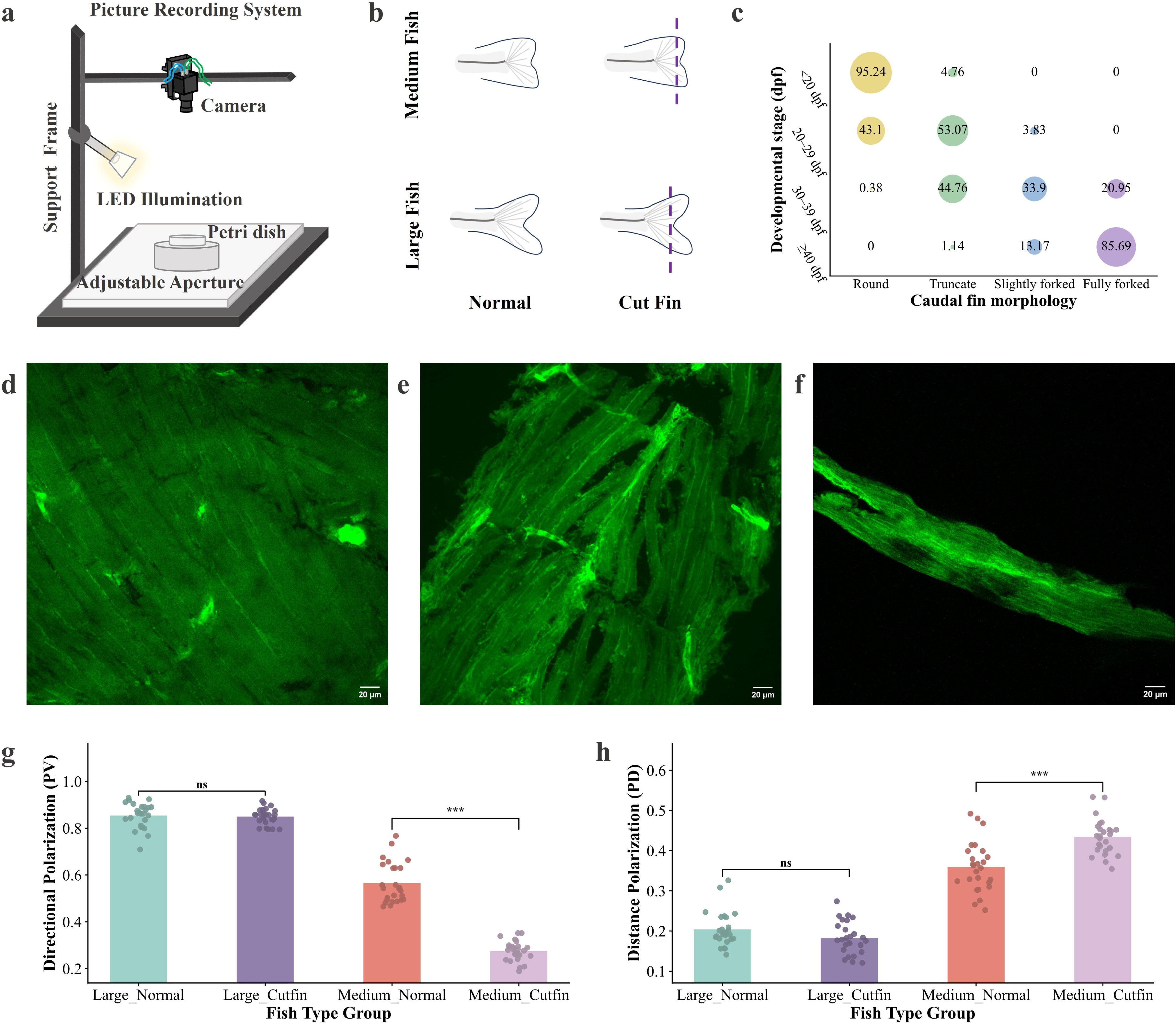
Morphological basis of the zebrafish locomotor system and its impact on group coordination. **a,** Schematic of the imaging setup for zebrafish morphology experiments, showing camera, supplementary light source, aperture adjustment module, support system, and experimental dish. **b,** Schematic of caudal fin ablation experiments in medium and large fish. Left: intact caudal fin. Right: caudal fin after removal of the forked region along the dashed line. **c,** Supplementary statistics of caudal fin morphology distribution across developmental stages. The x-axis shows four fin types (round, truncate, slightly forked, fully forked), and the y-axis represents developmental stages. Bubble size indicates the proportion of each fin type at the corresponding stage, with percentages labeled inside bubbles. **d-f,** Confocal imaging of muscle transgenic line CZ26 frozen sections at 40× magnification, with green fluorescence labeling muscle fiber structure. (d) Large fish. (e) Medium fish. (f) Small fish. **g-h,** Effect of caudal fin ablation on group coordination metrics. (g) Group directional polarization index (PV). (h) Group positional dispersion index (PD). Each dot represents one experimental video. Statistical comparisons between intact and fin-ablated groups were performed separately at the medium and large developmental stages using Welch’s t-test.

**Extended Data Figure 3.**
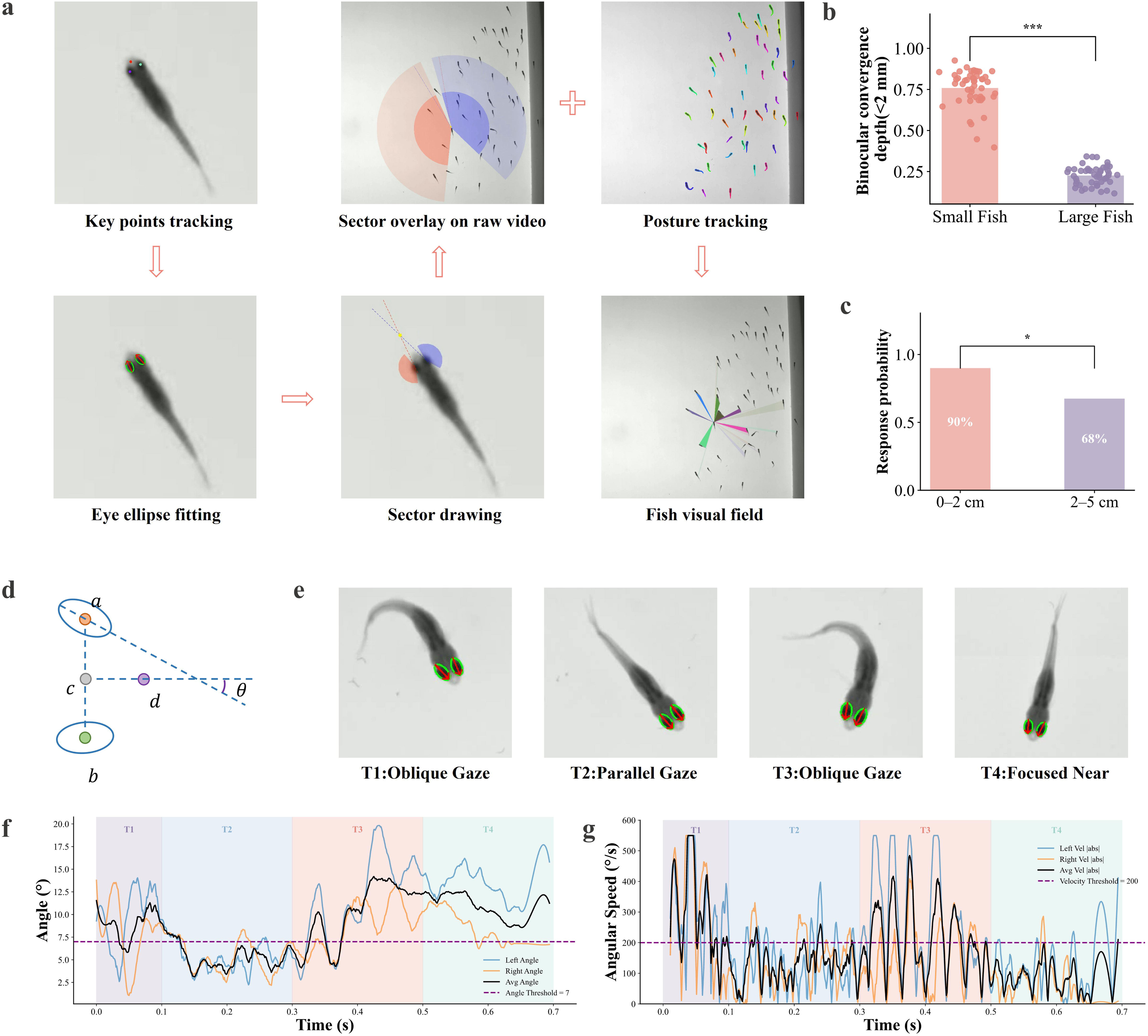
Eye-tracking and construction of the real visual perception model in zebrafish. **a,** Schematic of the workflow for eye-tracking and real visual perception modeling based on DeepLabCut [50]. Three key points on the left eye, right eye, and head are automatically tracked, and ellipses are fitted to each eye based on spatial positions to estimate eye-axis directions. Visual field sectors for each eye are constructed according to eye-axis orientation, and binocular convergence position is calculated. The local visual geometry model is then mapped back onto the original video frames, and combined with individual posture and visual distance constraints, ray-casting is used to determine truly perceivable neighbors, excluding occluded individuals, to construct an interaction network under real visual perception constraints. **b,** Comparison of the proportion of events with binocular convergence depth < 2 mm between small and large fish. Each dot represents one individual fish. Statistical comparisons were performed using Welch’s t-test. **c,** Probability of behavioral response in small fish to near (0-2 cm) versus medium (2-5 cm) light-spot stimulus in the light-spot experiment. Each data point represents the proportion of responding fish (n = 60 per stage). Statistical comparisons were performed using a two-proportion Z-test; asterisks indicate significance levels. **d,** Definition of eye movement angle. The head midline is determined by the midpoint between left and right eyes and the head key point; the angle θ between the unilateral eye major axis extension and the body midline is defined as eye movement amplitude. **e-g,** Representative single large-fish eye movement sequence recorded at 1000 fps for determining analysis thresholds. (e) Video frame snapshots showing four typical stages: T1 (Oblique gaze), T2 (Parallel gaze), T3 (Oblique gaze), and T4 (Focused near). T1 and T3 correspond to posture adjustment phases, T2 and T4 correspond to forward gliding, with T4 representing clear near-focus state. (f) Temporal change of binocular angle θ. Blue and orange lines indicate left and right eyes, black line represents mean, dashed line indicates threshold *θ* = 7°. (g) Temporal change of binocular angular velocity; dashed line indicates threshold 200°/*s*. These two thresholds are used to define large-amplitude eye movement events.

**Extended Data Figure 4.**
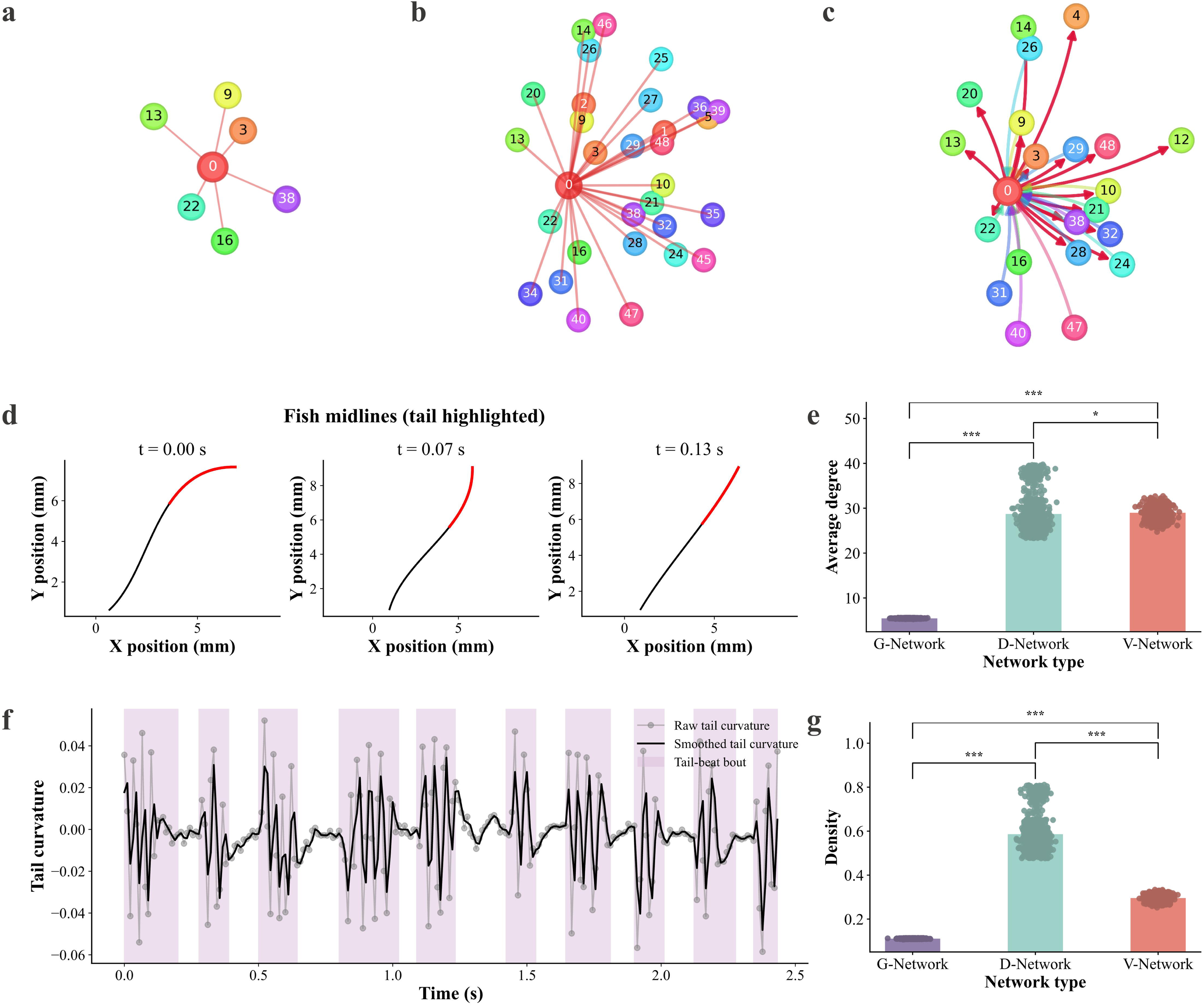
Local interaction networks under visual constraints and tail-beat dynamics. **a-c,** Schematic of local network structures constructed under three different interaction rules. To clearly illustrate network structure, only local subnetworks from the full 50-fish interaction network are shown. (a) Geometric neighborhood network (G-network) based on Delaunay triangulation. (b) Distance network (D-network) constructed using experimentally measured visual perceptual distance constraints. (c) Visual network (V-network) incorporating real visual perception and individual posture occlusion constraints. **d, f,** Workflow for extracting tail-beat dynamics. (d) Representative illustration of zebrafish midline fitting; the tail region is marked in red, showing midline morphology across three consecutive frames. (f) Example curve of tail curvature over time for a single fish. Gray dots represent raw data, black curve shows smoothed fit, and purple shaded area indicates tail-beat events. **e, g,** Quantitative comparison of network topological metrics under different network constructions. (e) Average degree. (g) Network density. Each dot represents one frame from the same video. Network metrics were computed for each frame using three different network construction methods. Pairwise comparisons of metric distributions across frames between methods were performed using Welch’s t-test.

**Extended Data Figure 5.**
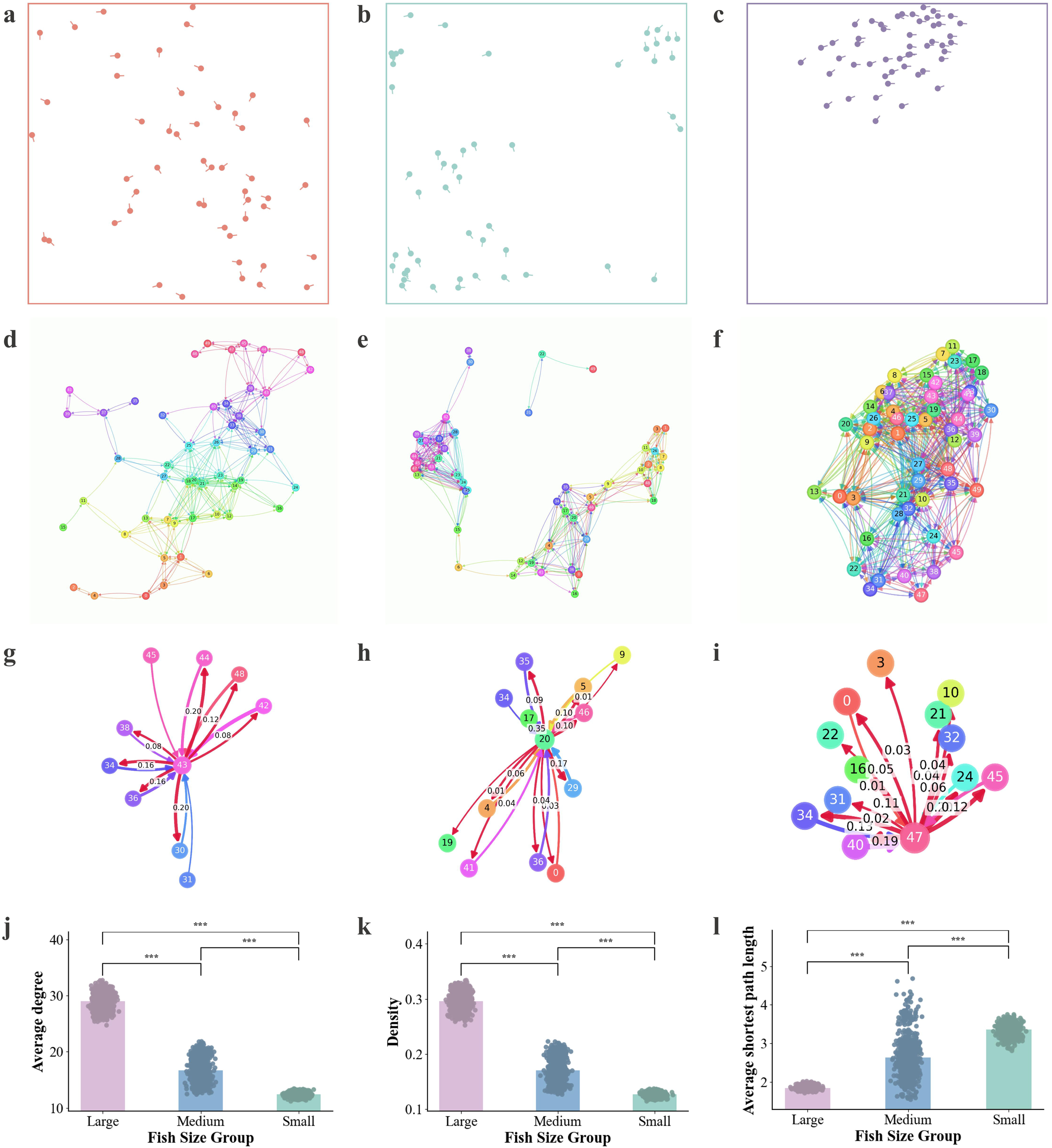
Group spatial structure and weighted interaction analysis based on the visual network. **a-c,** Spatial distribution and movement direction of 50 zebrafish at small, medium, and large fish stages. Dots indicate head positions, and short lines indicate instantaneous swimming direction. **d-f,** Illustration of group interaction structure based on the visual network (V-network) for small fish (d), medium fish (e), and large fish (f). **g-i,** Local structure of the weighted visual interaction network (W-network) constructed by incorporating distance decay and visual contour proportion on top of the V-network. A single fish is shown as the central node; red arrows indicate neighbors perceived by the central fish, and arrows of other colors indicate visual connections from other fish toward the central fish. Only the weights of connections from the central fish to its neighbors are labeled. Corresponding panels show small fish (g), medium fish (h), and large fish (i). **j-l,** Quantitative comparison of group network topology metrics based on the V-network across developmental stages. (j) Average degree; (k) Network Density; (l) Average shortest path length. Each dot represents one frame from the corresponding developmental-stage video. Network metrics were calculated for each frame, and distributions across frames were compared between developmental stages using Welch’s t-test.

**Extended Data Figure 6.**
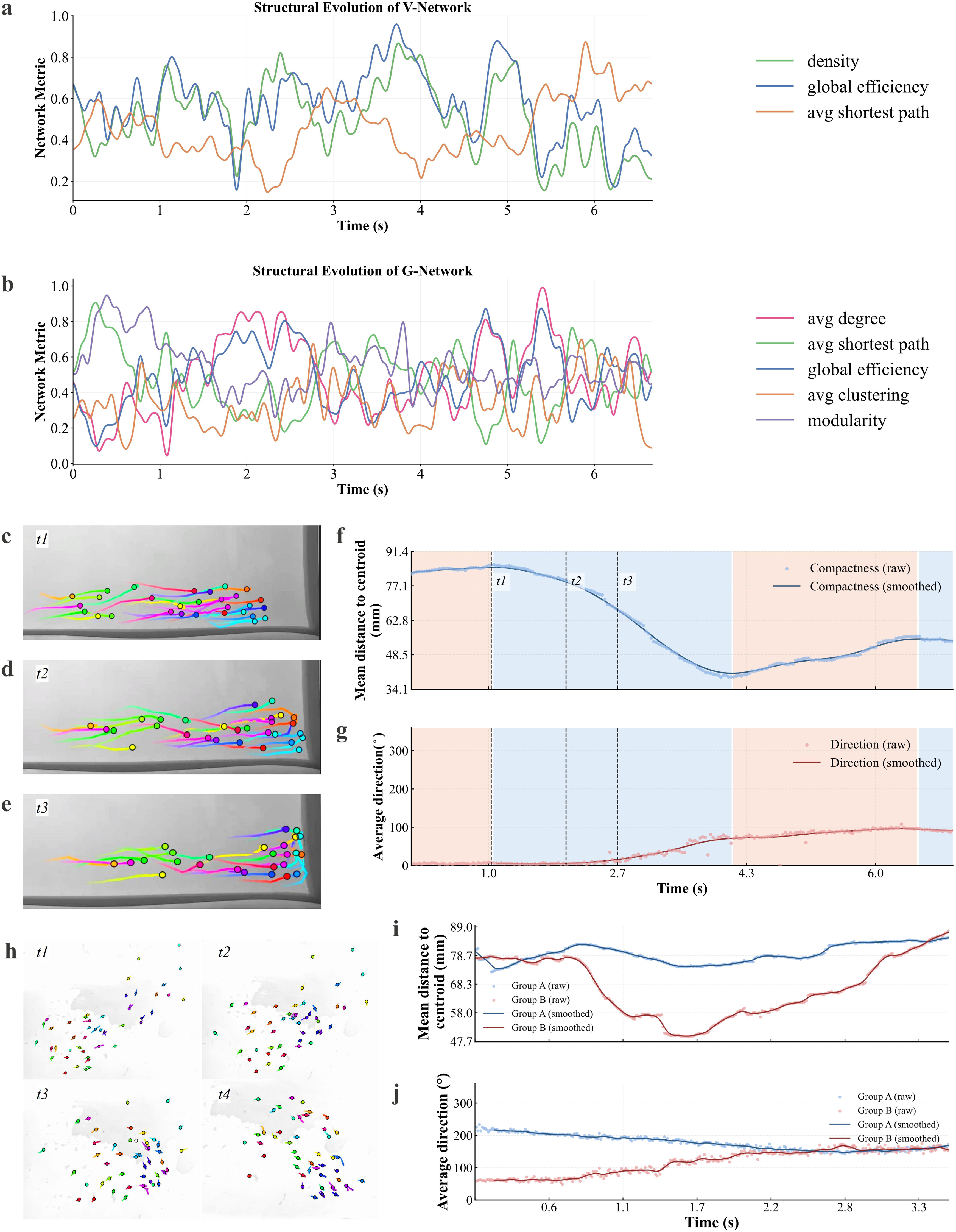
Network dynamics and group reorganization during turning under environmental constraints. **a,** Normalized group physical and network metrics over time based on the visual network (V-network), including individual density, global efficiency, and average shortest path length. **b,** Normalized topological metrics over time based on the geometric neighborhood network (G-network), including mean degree, average shortest path length, global efficiency, average clustering coefficient, and modularity. (C-E) Representative moments during group turning in a rectangular tank. **c,** Pre-turn stage (t1); **d,** Fish group approaching the corner region (t2); **e,** Onset of overall directional change (t3). **f-g,** Temporal evolution of group spatial structure and movement state during turning. **f,** Mean distance of individuals to the group centroid; **g,** Mean group movement direction. Scatter points represent experimental data, curves show fitted results. Shaded areas indicate contraction and expansion of the group envelope, and dashed lines correspond to key time points t1, t2, and t3. **h-j,** Representative process of merging two independent groups under environmental constraints. (h) Key time points t1-t4; (i) Mean distance of individuals to their respective group centroids; (j) Average group movement direction. Blue indicates group A, red indicates group B.

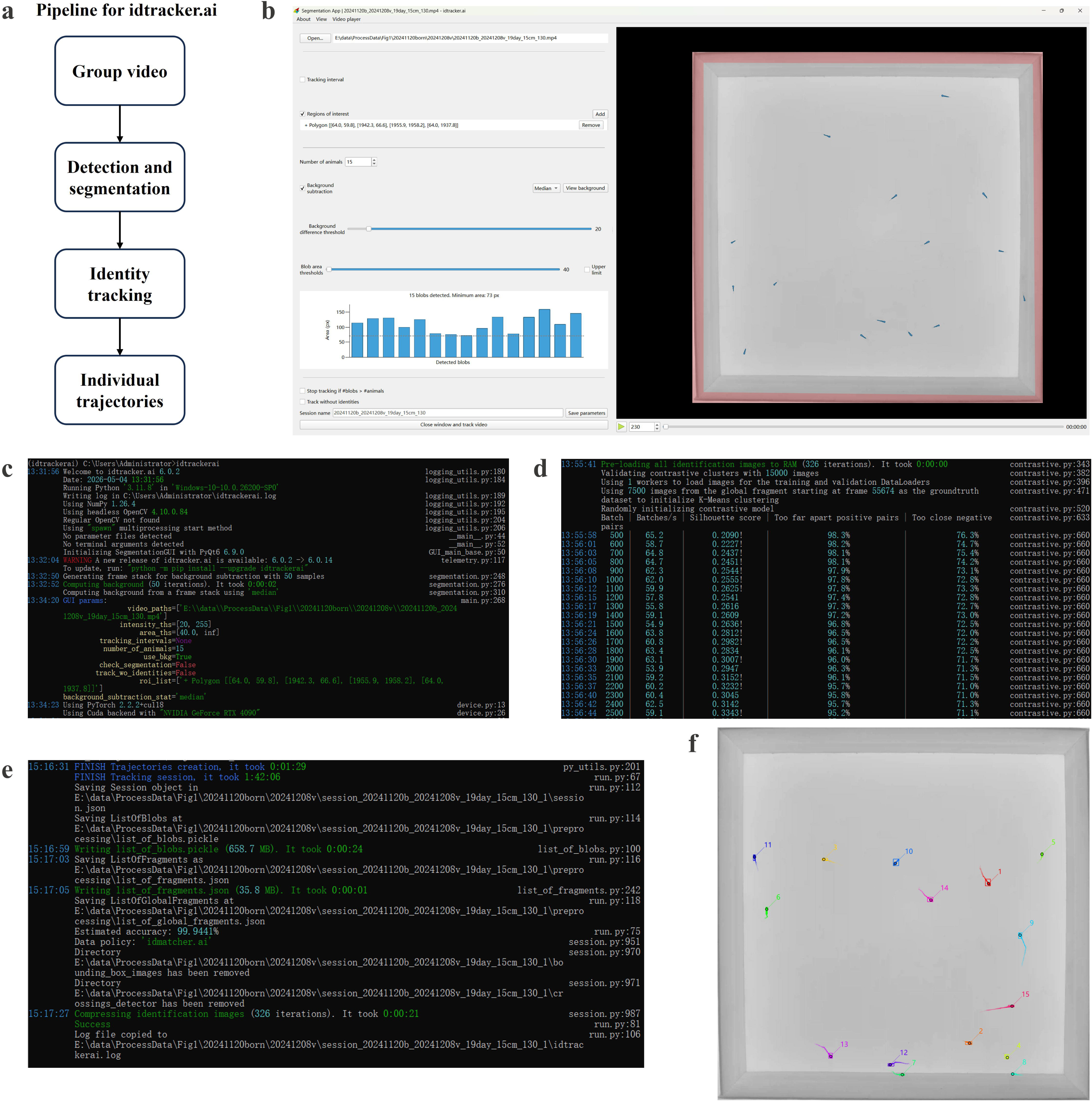

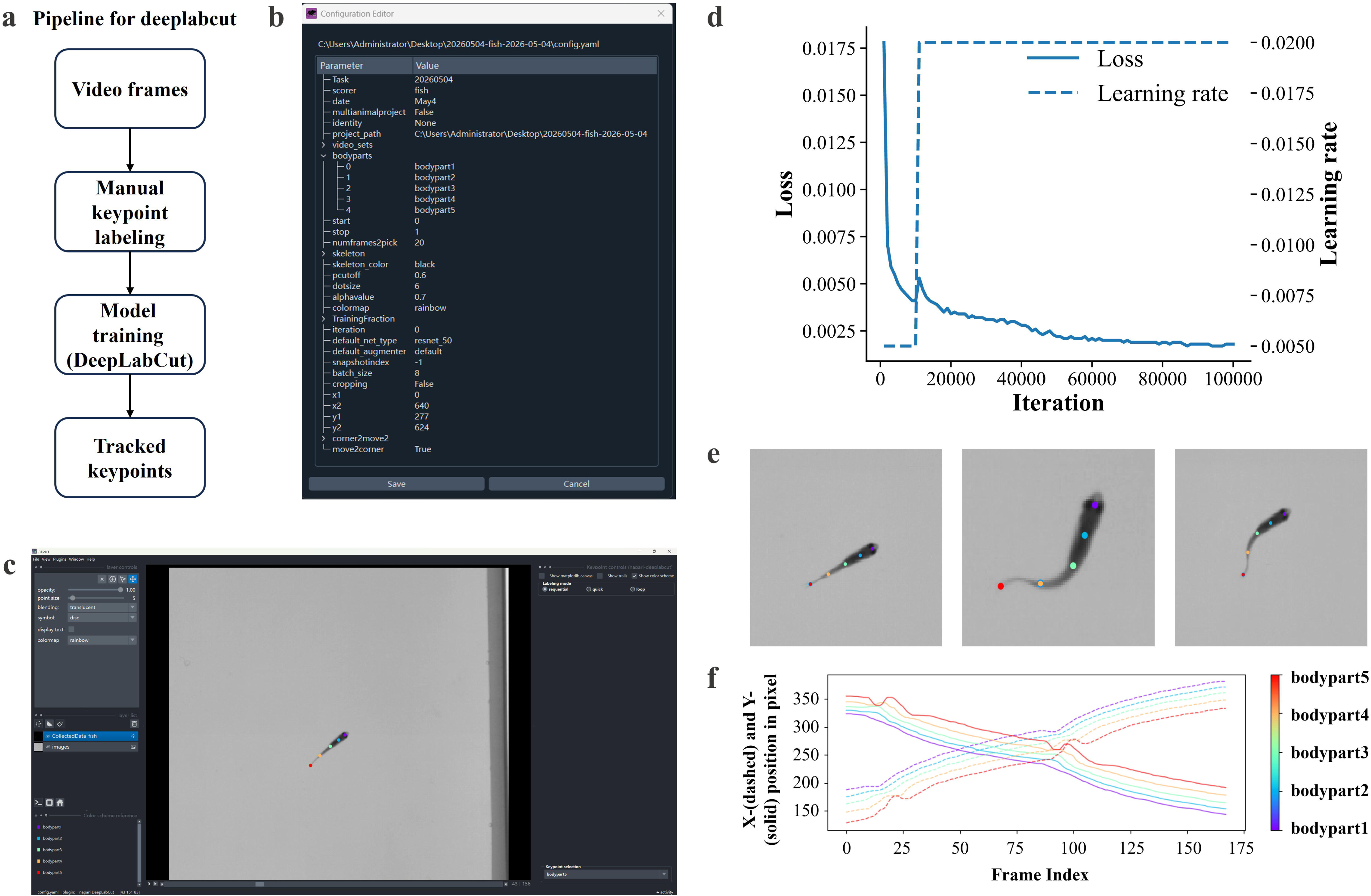

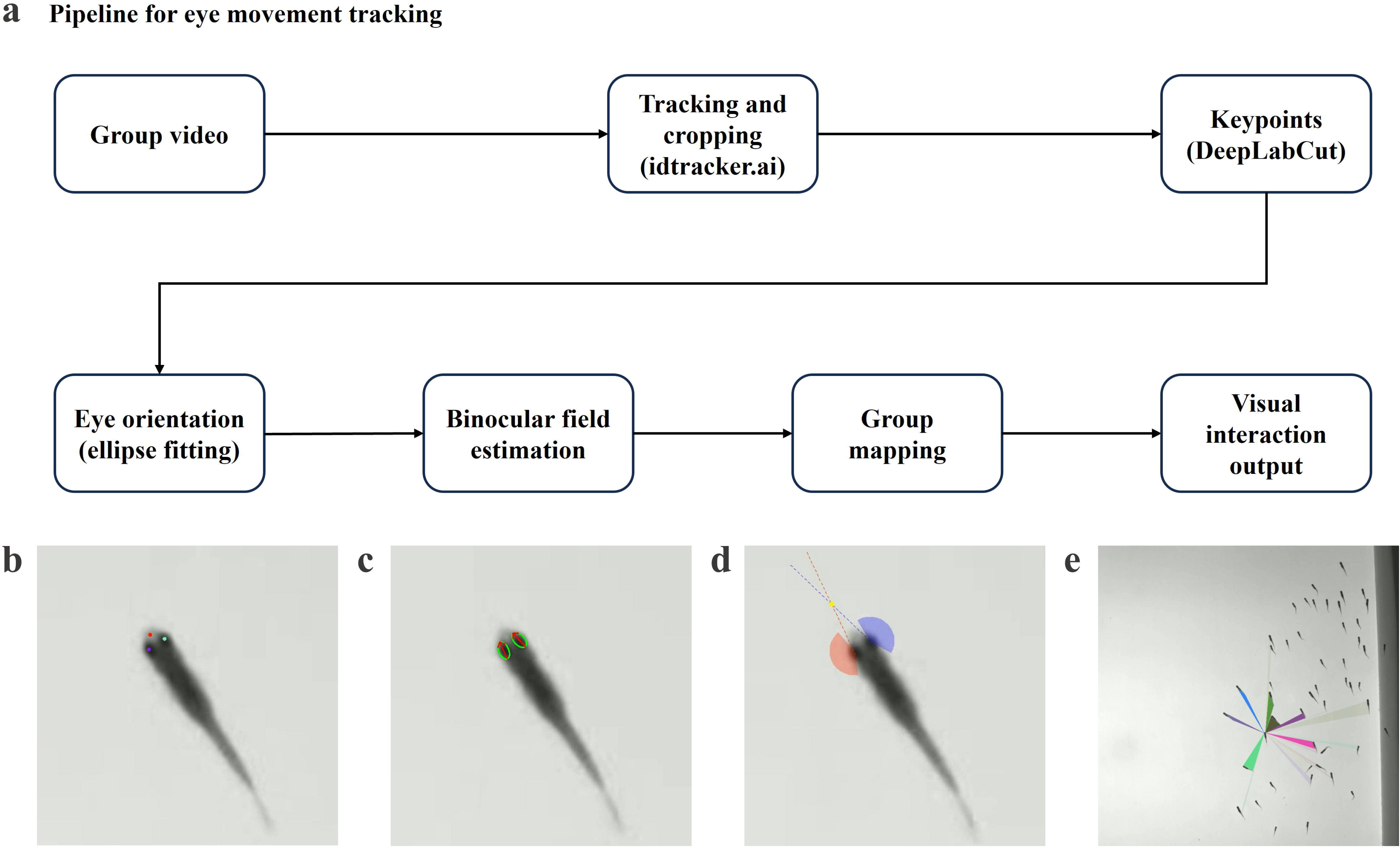

## Notes

### Competing Interest Statement

The authors have declared no competing interest.

