## Supplementary Information for "Origin of Schooling and Collective Environmental Adaptation in Zebrafish"

**This PDF file includes:**

Supplementary text

SI References

Legends for Movies S1 to S6

Caption for Figure S1 to S3

Figure S1 to S3

**Other supplementary materials for this manuscript include the following:**

Movies S1 to S6

**I. Multi-animal tracking using idtracker.ai**

Individual tracking and contour-based posture extraction were performed using idtracker.ai **(Figure S1)**. Group videos were processed within the idtracker.ai pipeline, where animals were first detected and segmented from the background on a frame-by-frame basis. To improve segmentation robustness, a region of interest (ROI) was manually defined to restrict analysis to the experimental arena and exclude irrelevant background regions.

Segmentation quality was further optimized by adjusting parameters such as the background difference threshold and blob area thresholds according to image contrast, illumination conditions, and fish size. These parameters were tuned to ensure reliable detection of individual animals while minimizing noise, fragmented detections, and false positives **(Figure S1b)**. The total number of individuals present in each recording was specified prior to tracking, providing a constraint for consistent identity assignment across frames. Following segmentation, idtracker.ai performed identity tracking by associating detections across frames and assigning a unique, persistent identity to each individual. The training and identification procedure was monitored during execution to ensure stable identity assignment and convergence of the tracking process **(Figure S1c-e)**. In cases of ambiguous detections or temporary occlusions, the algorithm maintained identity continuity based on learned visual features and temporal consistency.

Tracking results were subsequently inspected and validated using the built-in visualization tools **(Figure S1f)**, which allow simultaneous visualization of individual identities, body contours, and trajectories. This step enabled the identification of potential tracking errors, such as identity swaps or missed detections. These errors were minimized through iterative parameter refinement and, when necessary, manual correction of erroneous assignments to ensure consistency and accuracy of identity tracking. The resulting dataset consisted of continuous trajectories and contour information for each individual fish, exported as time-resolved positional coordinates (e.g., tabular or array formats). These data enabled direct reconstruction, visualization, and quantitative analysis of individual trajectories.

**II. Keypoint tracking using DeepLabCut**

Body posture and anatomical landmarks were quantified using DeepLabCut **(Figure S2)**. In this study, DeepLabCut was used to annotate multiple sets of body keypoints, including points along the body midline as well as the left eye, right eye, and head position. The general workflow for keypoint annotation and model training is illustrated in **(Figure S2a)**, with project configuration and keypoint definitions shown in Fig. S2b. Midline keypoints are presented as a representative example.

For each dataset, a project was created and configured to define the set of keypoints to be tracked **(Figure S2b)**. A subset of video frames was then sampled and manually annotated to generate the training dataset **(Figure S2c)**. Typically, several hundred frames were selected to ensure sufficient coverage of different body postures, orientations, and imaging conditions. These annotations were used to train a deep neural network within the DeepLabCut framework **(Figure S2d)**, typically using a ResNet-50 backbone and trained for approximately 100,000 iterations, providing a balance between accuracy and computational efficiency.

Following training, the network was applied to all frames to predict keypoint positions automatically. The tracking performance was evaluated by visual inspection of labeled videos **(Figure S2e)**, allowing assessment of spatial accuracy and temporal consistency. In cases where predictions were unreliable (e.g., due to occlusions, motion blur, or unusual postures), additional frames were incorporated into the training set and the model was iteratively refined to improve performance.

The final output consisted of time-resolved coordinates of all annotated keypoints for each individual, typically represented in pixel space **(Figure S2f)**. These data enabled reconstruction of body posture over time and served as the basis for subsequent analyses of eye movement, orientation, and visually guided interactions. To ensure consistency across individuals and recordings, the same keypoint definition and labeling scheme were applied across all datasets.

**III. Eye movement tracking**

Eye movement tracking was performed by integrating individual-level trajectory information with keypoint-based pose estimation **(Figure S3a)**. Based on previously obtained trajectories, local video regions centered on each individual were extracted to enable high-resolution analysis at the single-fish level. Keypoints corresponding to the left eye, right eye, and head position were then used to localize the eye region and provide a reference frame for orientation estimation **(Figure S3b)**.

Eye orientation was estimated using an ellipse-fitting approach combined with image-based processing. For each frame, a local region around the eye was extracted and converted to grayscale, followed by threshold-based segmentation to identify candidate eye regions. Elliptical contours were then fitted to these regions, and the major axis of each ellipse was taken as the principal orientation axis of the eye **(Figure S3c)**.

To ensure consistent interpretation, the orientation of the major axis was resolved relative to the head position, allowing the forward-facing direction to be determined unambiguously. Additional geometric constraints were applied to improve robustness, enabling reliable identification of left and right eyes under varying imaging conditions. Based on the orientations of the left and right eyes, binocular visual field sectors were constructed for each individual **(Figure S3d)**.

These visual features were subsequently mapped back to the original group coordinate system and integrated with multi-individual trajectories. For group-level analysis, a ray-casting approach was applied to determine which neighboring individuals fell within the effective visual field at each time point, excluding occluded targets **(Figure S3e)**. This enabled the construction of interaction networks under biologically realistic visual perception constraints.

**IV. Dynamical analysis.**

Directional polarization (PV) was defined as

$$\begin{aligned} \boldsymbol{PV=}\frac{\boldsymbol{1}}{\boldsymbol{N}}\left\| \sum_{\boldsymbol{i=1}}^{\boldsymbol{N}} \frac{\boldsymbol{v}_{\boldsymbol{i}}}{\left\| \boldsymbol{v}_{\boldsymbol{i}} \right\|} \right\|\boldsymbol{\#}\mathbf{Eq.S1} \end{aligned}$$

Where $N$ is the number of fish and $v_{i}$ is the instantaneous velocity vector of individual $i$. Each velocity vector was normalized to unit length prior to summation, and $\left\| \cdot\right\|$ denotes the Euclidean norm. PV ranges from 0 to 1, with larger values indicating stronger alignment of movement directions across the group.

Distance polarization (PD) was defined as

$$\begin{aligned} \boldsymbol{PD=}\frac{\boldsymbol{2}}{\boldsymbol{N}\left( \boldsymbol{N-1} \right)\boldsymbol{L}}\sum_{\boldsymbol{i<j}} \left\| \boldsymbol{x}_{\boldsymbol{i}}\boldsymbol{-}\boldsymbol{x}_{\boldsymbol{j}} \right\| \#\mathbf{Eq.S2} \end{aligned}$$

Where $N$ is the number of fish and $L$ is the side length of the experimental arena, and $\left\| x_{i}-x_{j} \right\|$ denotes the Euclidean distance between fish $i$ and $j$. PD ranges from 0 to 1, with smaller values indicating higher spatial compactness of the group.

Average speed **(Extended Data Fig. 1H)** and average acceleration **(Extended Data Fig. 1I).** They were used to characterize overall swimming intensity and dynamic variability of the group. These metrics were computed within time windows selected based on maximal directional polarization and provide complementary kinematic descriptions of collective organization.

Average speed was defined as

$$\begin{aligned} \overline{\boldsymbol{s}}\boldsymbol{=}\left\langle\left\| \boldsymbol{v}_{\boldsymbol{i}}\left( \boldsymbol{t} \right) \right\| \right\rangle_{\boldsymbol{i,t}}\boldsymbol{\#}\mathbf{Eq.S3} \end{aligned}$$

Where $N$ is the number of fish, $v_{i}(t)$ denotes the instantaneous velocity vector of individual $i$ at time $t$, and $\left\langle\cdot\right\rangle_{i,t}$ represents averaging over all individuals and frames. Larger values indicate stronger overall swimming activity.

Average acceleration was defined as

$$\begin{aligned} \overline{\boldsymbol{a}}\boldsymbol{=}\left\langle\left\| \frac{\boldsymbol{v}_{\boldsymbol{i}}\left( \boldsymbol{t+1} \right)\boldsymbol{-}\boldsymbol{v}_{\boldsymbol{i}}\left( \boldsymbol{t} \right)}{\boldsymbol{\Delta t}} \right\| \right\rangle_{\boldsymbol{i,t}}\boldsymbol{\#}\mathbf{Eq.S4} \end{aligned}$$

Where $\Delta t$ is the inter-frame time interval. Average acceleration reflects the magnitude of velocity fluctuations and quantifies the dynamic activity and stability of group motion.

Tail-beat synchrony **(Fig. 4e-g).** It was introduced to quantify local coordination of tail movements within the group. Neighbor relationships were determined frame by frame using the directed visibility network (V-network), in which nodes represent individuals and edges indicate visual connections.

Let $b_{i}(t)\in\{0,1\}$ denote the tail-beat state of fish $i$ at time $t$, where $b_{i}(t)=1$ indicates a tail beat and $b_{i}(t)=0$ indicates no tail beat. Let $N_{i}(t)$ denote the set of outgoing neighbors of fish $i$ in the V-network at time $t$.

Neighborhood tail-beat synchrony was defined as

$$\begin{aligned} \boldsymbol{S}_{\boldsymbol{i}}\left( \boldsymbol{t} \right)\boldsymbol{=}\frac{\boldsymbol{1}}{\left| \boldsymbol{N}_{\boldsymbol{i}}\left( \boldsymbol{t} \right) \right|}\sum_{\boldsymbol{j\in}\boldsymbol{N}_{\boldsymbol{i}}\left( \boldsymbol{t} \right)} \boldsymbol{b}_{\boldsymbol{j}}\left( \boldsymbol{t} \right)\boldsymbol{\#}\mathbf{Eq.S5} \end{aligned}$$

where $\left| N_{i}(t) \right|$ is the number of neighbors of fish $i$. This metric represents the fraction of neighbors exhibiting simultaneous tail beats when fish $i$ performs a tail beat. Synchrony was computed only when $b_{i}(t)=1$ and $\left| N_{i}(t) \right|>0$; otherwise, values were treated as missing. The resulting frame-by-individual synchrony matrix was used for visualization and subsequent statistical analysis. $S_{i}(t)$ ranges from 0 to 1, with larger values indicating stronger local tail-beat coordination.

Mean group distance and mean movement direction **(Fig. 4i-j, Fig. 4l-m, Extended Data Fig. 6f-g, Extended Data Fig. 6i-j).** They were computed to characterize temporal changes in spatial contraction and collective turning behavior.

At each frame $t$, let $x_{i}(t)$ denote the position of fish $i$, and $N_{v}(t)$ the number of valid individuals. The group centroid was defined as

$$\begin{aligned} \boldsymbol{c}\left( \boldsymbol{t} \right)\boldsymbol{=}\frac{\boldsymbol{1}}{\boldsymbol{N}_{\boldsymbol{v}}\left( \boldsymbol{t} \right)}\sum_{\boldsymbol{i=1}}^{\boldsymbol{N}_{\boldsymbol{v}}\left( \boldsymbol{t} \right)} \boldsymbol{x}_{\boldsymbol{i}}\left( \boldsymbol{t} \right)\boldsymbol{\#}\mathbf{Eq.S6} \end{aligned}$$

Mean group distance was defined as

$$\begin{aligned} \boldsymbol{D}\left( \boldsymbol{t} \right)\boldsymbol{=}\frac{\boldsymbol{1}}{\boldsymbol{N}_{\boldsymbol{v}}\left( \boldsymbol{t} \right)}\sum_{\boldsymbol{i=1}}^{\boldsymbol{N}_{\boldsymbol{v}}\left( \boldsymbol{t} \right)} \left\| \boldsymbol{x}_{\boldsymbol{i}}\left( \boldsymbol{t} \right)\boldsymbol{-c}\left( \boldsymbol{t} \right) \right\|\boldsymbol{\#}\mathbf{Eq.S7} \end{aligned}$$

where $\parallel\cdot\parallel$ denotes the Euclidean norm. Smaller values of $D(t)$ indicate stronger spatial aggregation of the group.

The instantaneous velocity of fish $i$ was computed as $v_{i}(t)=x_{i}(t)-x_{i}(t-1)$, and normalized to unit length prior to averaging. The mean group direction vector was defined as

$$\begin{aligned} \boldsymbol{V}\left( \boldsymbol{t} \right)\boldsymbol{=}\frac{\boldsymbol{1}}{\boldsymbol{N}_{\boldsymbol{v}}\left( \boldsymbol{t} \right)}\sum_{\boldsymbol{i=1}}^{\boldsymbol{N}_{\boldsymbol{v}}\left( \boldsymbol{t} \right)} \frac{\boldsymbol{v}_{\boldsymbol{i}}\left( \boldsymbol{t} \right)}{\left\| \boldsymbol{v}_{\boldsymbol{i}}\left( \boldsymbol{t} \right) \right\|}\boldsymbol{\#}\mathbf{Eq.S8} \end{aligned}$$

The mean movement direction of the group was obtained as the polar angle of $V(t)$:

$$\begin{aligned} \boldsymbol{\theta}\left( \boldsymbol{t} \right)\boldsymbol{=atan}\boldsymbol{2}\left( \boldsymbol{V}_{\boldsymbol{y}}\left( \boldsymbol{t} \right)\boldsymbol{,}\boldsymbol{V}_{\boldsymbol{x}}\left( \boldsymbol{t} \right) \right)\boldsymbol{\#}\mathbf{Eq.S9}\boldsymbol{\#} \end{aligned}$$

where $V_{x}(t)$and $V_{y}(t)$are the Cartesian components of $V(t)$in the experimental coordinate system. Angle values were converted to the range $0^{\circ}-360^{\circ}$.

**III. Network construction and quantification.**

W-Network. The V-network is a directed graph, where nodes represent individuals and edges indicate visibility from the observer to the target. For each frame, the Euclidean distance between individual $i$(observer) and $j$(target) was computed as

$$\begin{aligned} \boldsymbol{d}_{\boldsymbol{ij}}\boldsymbol{=}\sqrt{\left( \boldsymbol{x}_{\boldsymbol{i}}\boldsymbol{-}\boldsymbol{x}_{\boldsymbol{j}} \right)^{\boldsymbol{2}}\boldsymbol{+}\left( \boldsymbol{y}_{\boldsymbol{i}}\boldsymbol{-}\boldsymbol{y}_{\boldsymbol{j}} \right)^{\boldsymbol{2}}}\boldsymbol{\#}\mathbf{Eq.S10} \end{aligned}$$

Only pairs satisfying $d_{ij}\leq d_{c}$ were considered as potential visual connections, where the visual threshold $d_{c}$ was determined from light-spot stimulation experiments and scaled by body length.

Occlusion was evaluated using a ray-casting method based on eye-tracking data **(Extended Data Fig. 3A)**. Discrete rays were emitted from the binocular viewing directions, and intersections with body contours were detected. A directed edge $i\to j$ was established only if the target individual lay within the observer’s visual field and was not fully occluded.

On this basis, a weighted vision network [1] (W-Network) was further constructed. For each visual edge $i\to j$, the visual angle $\theta_{ij}$ subtended by the target and the mean distance $\overline{d}_{ij}$ were computed. A distance-dependent factor $f(\overline{d}_{ij})$, obtained from behavioral response probabilities in the light-spot experiments, was applied. The visual weight was defined as

$$\begin{aligned} \boldsymbol{w}_{\boldsymbol{ij}}\boldsymbol{=}\frac{\boldsymbol{\theta}_{\boldsymbol{ij}}\boldsymbol{f}\left( {\overline{\boldsymbol{d}}}_{\boldsymbol{ij}} \right)}{\sum_{\boldsymbol{k}} \boldsymbol{\theta}_{\boldsymbol{ik}}\boldsymbol{f}\left( {\overline{\boldsymbol{d}}}_{\boldsymbol{ij}} \right)}\boldsymbol{\#}\mathbf{Eq.S11}\boldsymbol{\#}\boldsymbol{\#} \end{aligned}$$

Where the denominator sums over all targets visible to observer $i$, ensuring normalization of outgoing weights. This framework integrates visual range, directional field of view, occlusion, and distance-dependent sensitivity, yielding a weighted directed network that approximates realistic visual interactions within the group.

Average degree. The average degree [2-5] was defined as

$$\begin{aligned} \left\langle\boldsymbol{K} \right\rangle\boldsymbol{=}\frac{\boldsymbol{2}\boldsymbol{E}}{\boldsymbol{N}}\boldsymbol{\#}\mathbf{Eq.S12} \end{aligned}$$

where $E$ is the total number of edges and $N$ is the number of nodes in the network. This definition applies to both undirected and directed networks; for directed networks, in-degree and out-degree were combined. The average degree quantifies the mean number of interaction links per individual and reflects the overall richness of visual connections within the group.

Network density. Network density [2-5] was defined as the ratio between the number of existing edges and the maximum possible number of edges.

For directed networks (without self-loops), density was computed as

$$\begin{aligned} \boldsymbol{D=}\frac{\boldsymbol{E}}{\boldsymbol{N}\left( \boldsymbol{N-1} \right)}\boldsymbol{\#}\mathbf{Eq.S13} \end{aligned}$$

Where $E$ is the number of directed edges and $N$ is the number of nodes.

For undirected networks, density was defined as

$$\begin{aligned} \boldsymbol{D=}\frac{\boldsymbol{2}\boldsymbol{E}}{\boldsymbol{N}\left( \boldsymbol{N-1} \right)}\boldsymbol{\#}\mathbf{Eq.S14} \end{aligned}$$

Network density quantifies the overall tightness of interactions within the group, with higher values indicating more densely connected individuals.

Average shortest path length [2-5]. Because the visual network is not necessarily strongly connected, shortest paths were computed only for node pairs connected by a directed path. Let $l_{ij}$ denote the shortest path length from node $i$ to node $j$. The average shortest path length was defined as

$$\begin{aligned} \boldsymbol{L=}\frac{\boldsymbol{1}}{\left| \boldsymbol{S} \right|}\sum_{\left( \boldsymbol{i,j} \right)\boldsymbol{\in S}} \boldsymbol{l}_{\boldsymbol{ij}}\boldsymbol{\#}\mathbf{Eq.S15} \end{aligned}$$

Where $S$ denotes the set of all reachable node pairs $(i,j)$. This metric reflects the average number of steps required for visual information to propagate across the group and was used to quantify the global communication efficiency of the network.

Global efficiency. Global efficiency [2-5] quantifies the overall efficiency of information transfer across the network and is defined as

$$\begin{aligned} \boldsymbol{E}_{\boldsymbol{glob}}\boldsymbol{=}\frac{\boldsymbol{1}}{\boldsymbol{N}\left( \boldsymbol{N-1} \right)}\sum_{\boldsymbol{i\neq j}} \frac{\boldsymbol{1}}{\boldsymbol{d}_{\boldsymbol{ij}}}\boldsymbol{\#}\mathbf{Eq.S16} \end{aligned}$$

Where $N$ is the number of nodes and $d_{ij}$ denotes the shortest path length from node $i$ to node $j$. For unreachable node pairs, $1/d_{ij}$ was set to 0. For undirected networks, $d_{ij}$ corresponds to the shortest undirected path length, whereas for directed networks only directed paths from $i$ to $j$ were considered. Compared with average shortest path length, global efficiency explicitly incorporates disconnected node pairs and therefore provides a more robust measure of network-wide communication efficiency.

Average clustering coefficient. The average clustering coefficient [2-5] quantifies the tendency of neighboring nodes to form local clusters and is defined as

$$\begin{aligned} \boldsymbol{C=}\frac{\boldsymbol{1}}{\boldsymbol{N}}\sum_{\boldsymbol{i=1}}^{\boldsymbol{N}} \frac{\boldsymbol{2}\boldsymbol{e}_{\boldsymbol{i}}}{\boldsymbol{k}_{\boldsymbol{i}}\left( \boldsymbol{k}_{\boldsymbol{i}}\boldsymbol{-1} \right)}\boldsymbol{\#}\mathbf{Eq.S17} \end{aligned}$$

Where $k_{i}$ is the degree of node $i$, and $e_{i}$ is the number of edges among its neighbors. A higher clustering coefficient indicates stronger local connectivity and tighter group structure.

Modularity. Modularity [2-5] measures the strength of community structure in the network and is defined as

$$\begin{aligned} \boldsymbol{Q=}\frac{\boldsymbol{1}}{\boldsymbol{2}\boldsymbol{E}}\sum_{\boldsymbol{i,j}} \left[ \boldsymbol{A}_{\boldsymbol{ij}}\boldsymbol{-}\frac{\boldsymbol{k}_{\boldsymbol{i}}\boldsymbol{k}_{\boldsymbol{j}}}{\boldsymbol{2}\boldsymbol{E}} \right]\boldsymbol{\delta}\left( \boldsymbol{c}_{\boldsymbol{i}}\boldsymbol{,}\boldsymbol{c}_{\boldsymbol{j}} \right)\boldsymbol{\#}\mathbf{Eq.S18} \end{aligned}$$

Where $A_{ij}$ denotes the adjacency matrix, $k_{i}$ is the degree of node node $i$, and $\delta\left( c_{i},c_{j} \right)=1$ if nodes $i$ and $j$ belong to the same community and 0 otherwise. Higher modularity values indicate more pronounced community separation within the group.

For environmental response analysis, all metrics were normalized and smoothed prior to visualization. Each time series $y$ was first min-max normalized as

$\begin{aligned} \boldsymbol{y}_{\boldsymbol{norm}}\boldsymbol{=}\frac{\boldsymbol{y-}\min\left( \boldsymbol{y} \right)}{\max\left( \boldsymbol{y} \right)\boldsymbol{-}\min\left( \boldsymbol{y} \right)}\boldsymbol{\#}\mathbf{Eq.S19} \end{aligned}$

**Legends for Supplementary Movies**

**Movie S1.** A representative video illustrating the analysis pipeline. The field of view is divided into three panels. Multi-animal pose tracking **(left)** shows each fish segmented with a green contour and represented by multiple color-coded keypoints along the body midline. Eye-tracking results **(middle)** show fitted elliptical contours of the eyes, associated angular sectors, and the intersection of the major axes used to estimate gaze direction. Visualization of the ego-centric V-network **(right)** shows a focal fish sampling its visual field; colored rays indicate the proportion of neighboring fish visible in different directions, taking occlusion into account.

**Movie S2.** Representative behavioral recordings of small **(left)**, medium **(middle)**, and large **(right)** fish. Small fish display largely independent swimming, medium fish aggregate into several small groups, and large fish form stable schooling with coherent group motion. The fish are colored by their normalized speed.

**Movie S3.** Representative recording of large fish following tail fin ablation. Fish still exhibit schooling behavior.

**Movie S4.** A representative recording showing the effect of light removal on group behavior. At the beginning, top illumination produces visible reflections on the water surface, indicating the light-on condition. When the light is switched off, the reflections disappear (infrared illumination only), and the organized group structure rapidly breaks down into disordered motion.

**Movie S5.** Representative examples of behavioral responses to light stimulation in large fish. Three distinct responses are shown: rapid avoidance **(left)**, active exploration toward the light source **(middle)**, and absence of an observable response **(right)**.

**Movie S6.** Visualization of different interaction networks for the same group of fish. The G-network **(left)**, D-network **(middle)**, and V-network **(right)** are shown.

**Caption for Supplementary Figures**

**Figure S1. Multi-animal tracking using idtracker.ai. a,** Overview of the idtracker.ai processing pipeline. **b,** Detection and segmentation interface used to define the region of interest, set the number of individuals, and adjust parameters for background subtraction and object detection. **c-e**, Representative stages of the tracking process **f,** Example output showing identity assignment, where different colors and IDs indicate individual zebrafish, together with extracted body contours and bounding boxes.

**Figure S2. Keypoint tracking using DeepLabCut. a,** Overview of the DeepLabCut workflow. **b,** Configuration of the project file, including definition of keypoints and training parameters. **c,** Manual annotation of selected video frames for training dataset generation. **d,** Training process of the neural network across iterations. **e,** Representative frames showing predicted keypoints overlaid on the fish body. **f,** Example output of keypoint coordinates in pixel space.

**Figure S3. Eye movement tracking and visual field reconstruction. a,** Overview of the eye movement tracking pipeline. **b,** Detection of left eye, right eye, and head keypoints. **c,** Ellipse fitting of eye regions, with fitted contours shown in green and the major axis (eye orientation) in red. **d,** Construction of binocular visual field sectors based on left and right eye orientations, with the convergence region indicating the estimated direction of visual attention. **e,** Visualization of visual fields at the group level using a ray-casting approach, showing which neighboring individuals fall within the effective visual range.

**Figure S1**

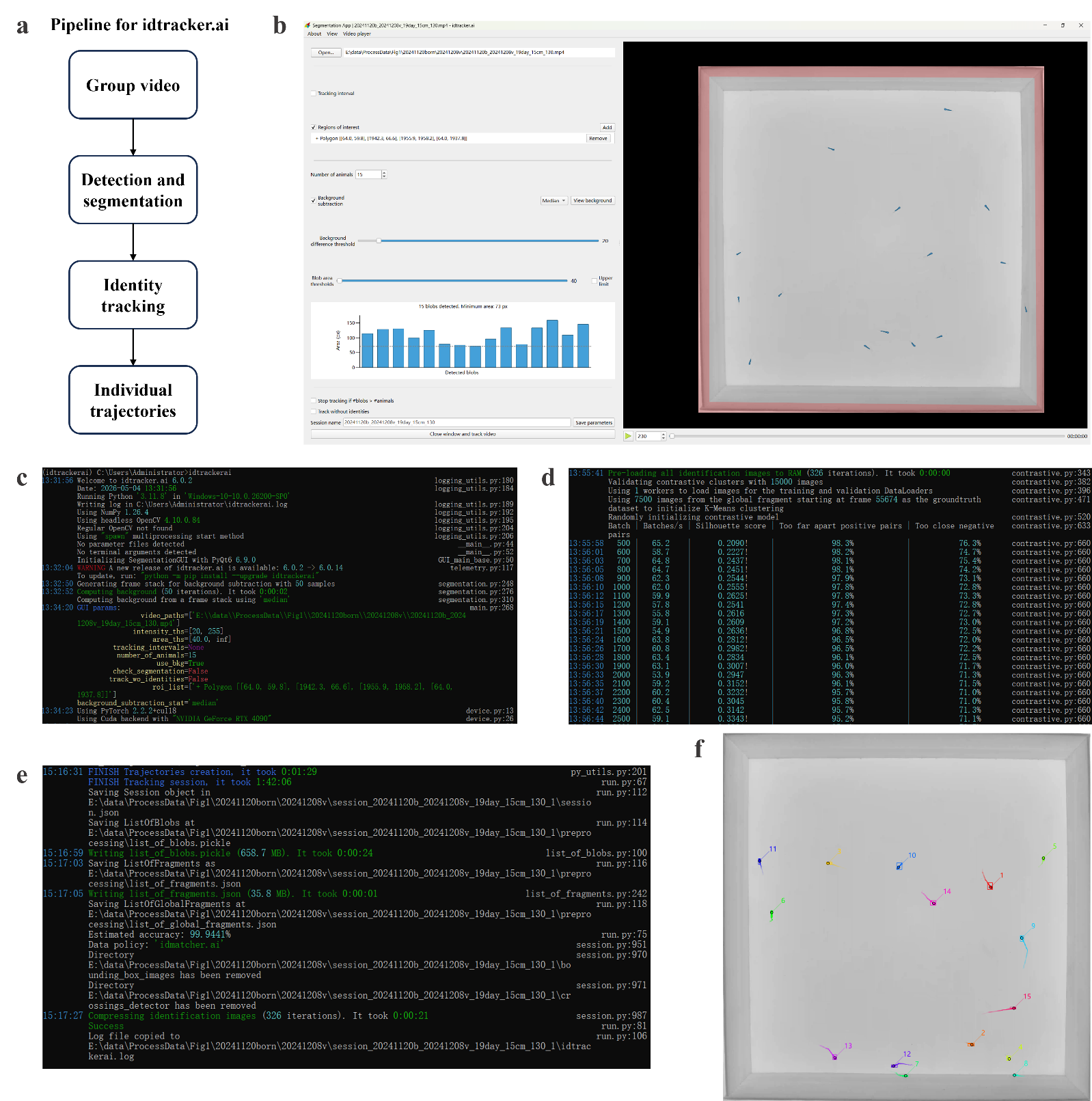

**Figure S2**

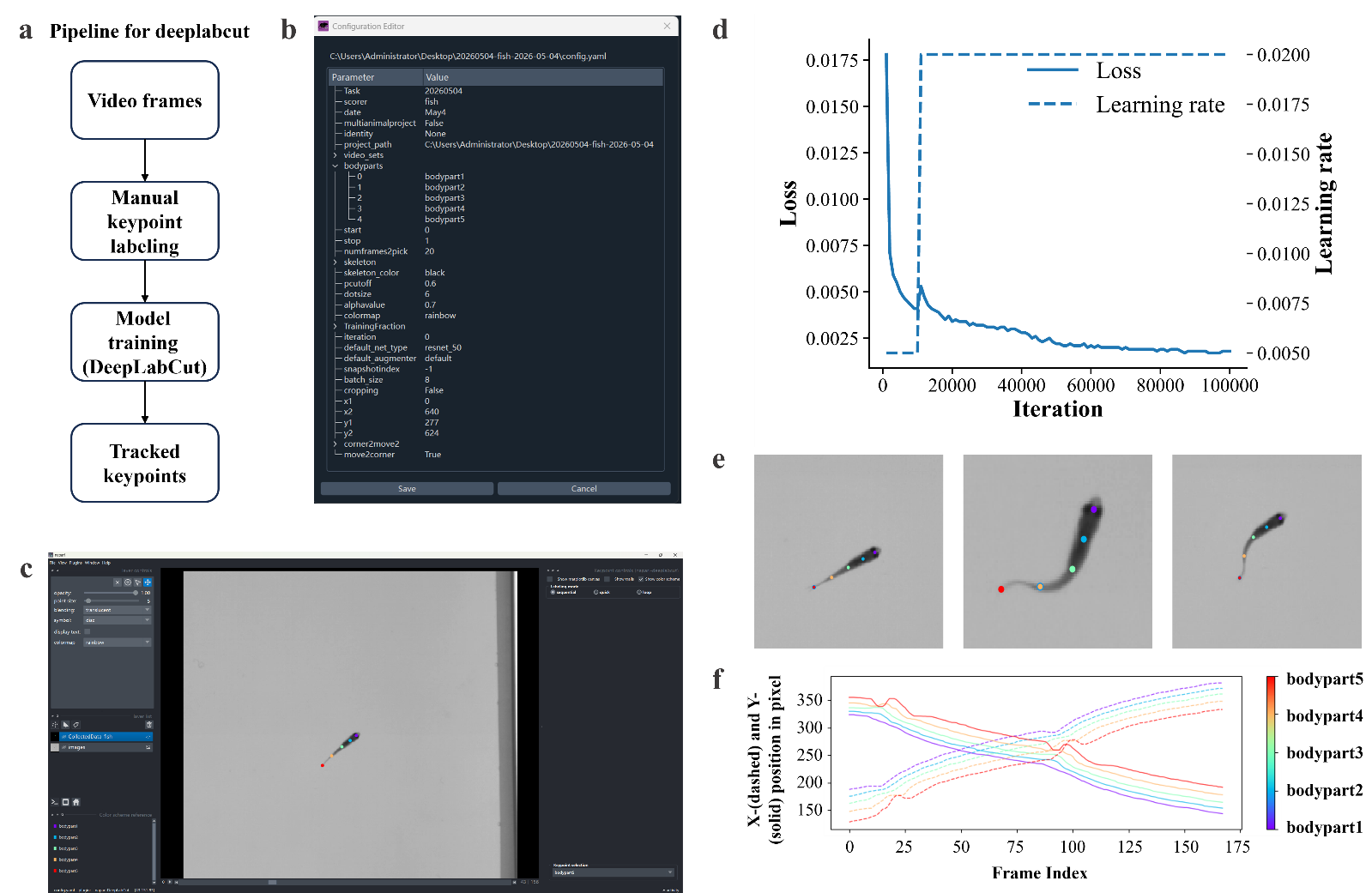

**Figure S3**

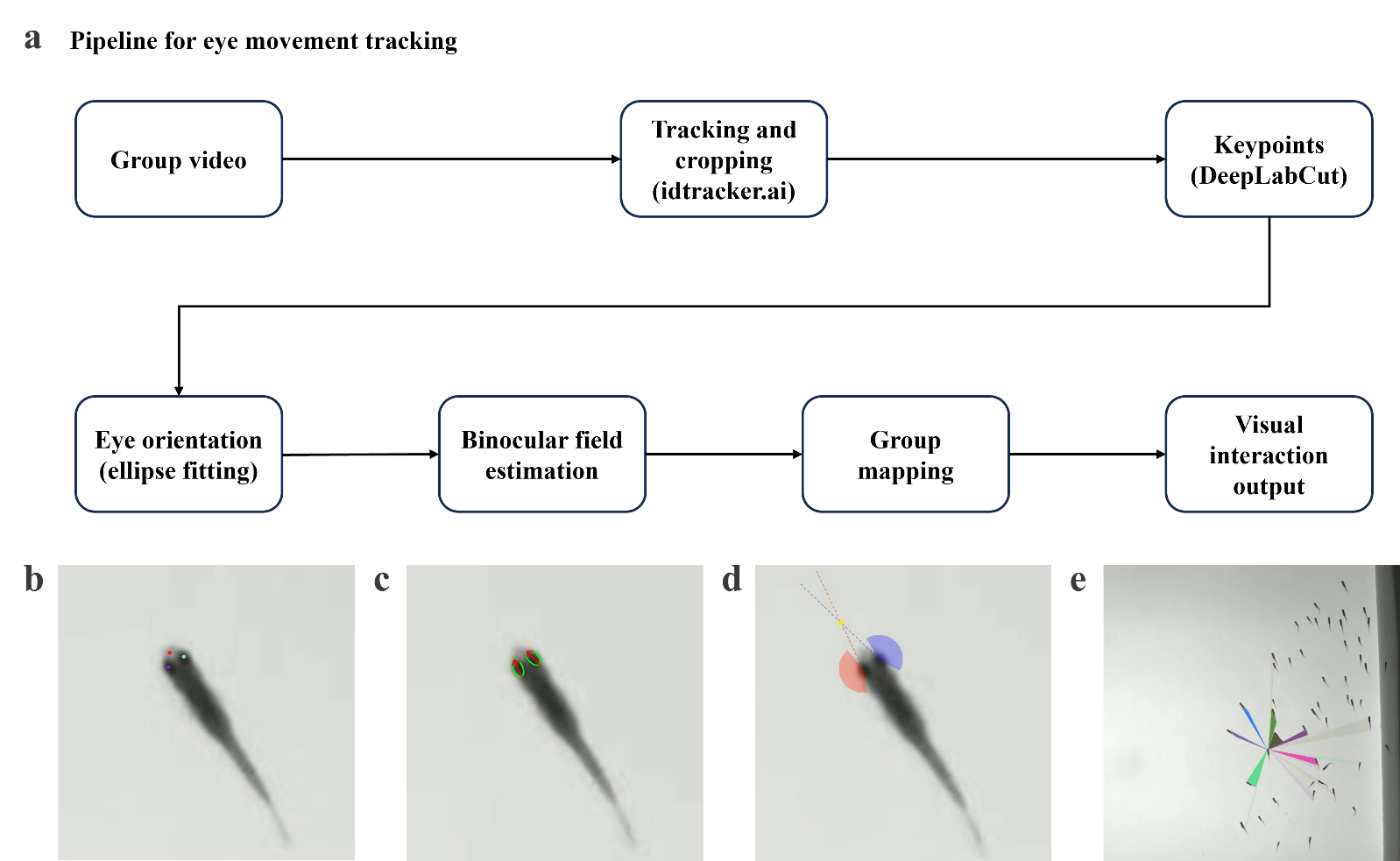
